# Neural dynamics at successive stages of the ventral visual stream are consistent with hierarchical error signals

**DOI:** 10.1101/092551

**Authors:** Elias B. Issa, Charles F. Cadieu, James J. DiCarlo

## Abstract

Ventral visual stream neural responses are dynamic, even for static image presentations. However, dynamical neural models of visual cortex are lacking as most progress has been made modeling static, time-averaged responses. Here, we studied population neural dynamics during face detection across three cortical processing stages. Remarkably, ~30 milliseconds after the initially evoked response, we found that neurons in intermediate level areas decreased their preference for faces, becoming anti-face preferring on average even while neurons in higher level areas achieved and maintained a face preference. This pattern of hierarchical neural dynamics was inconsistent with extensions of standard feedforward circuits that implemented recurrence within a cortical stage. Rather, recurrent models computing errors between stages captured the observed temporal signatures. Without additional parameter fitting, this model of neural dynamics, which simply augments the standard feedforward model of online vision to encode errors, also explained seemingly disparate dynamical phenomena in the ventral stream.

## INTRODUCTION

The primate ventral visual stream is a hierarchically organized set of cortical areas beginning with the primary visual cortex (V1) and culminating with distributed patterns of neural firing across the inferior temporal cortex (IT) that explicitly encode objects (i.e. linearly decodable object identity) (Hung, Kreiman, Poggio, & DiCarlo, 2005) and quantitatively account for core invariant object discrimination behavior in primates (Majaj, Hong, Solomon, & DiCarlo, 2015). Formalizing object recognition as the result of a series of feedforward computations yields models that achieve impressive performance in object categorization (Krizhevsky, Sutskever, & Hinton, 2012)(Zeiler & Fergus, 2013) similar to the absolute level of performance achieved by IT neural populations, and these models are the current best predictors of neural responses in IT cortex and its primary input layer, V4 (Cadieu et al., 2014)(Yamins et al., 2014). Thus, the feedforward inference perspective provides a simple but powerful, first-order framework for the ventral stream and core invariant object recognition.

However, visual object recognition behavior may not be executed via a single feedforward neural processing pass (a.k.a. feedforward inference) because IT neural responses are well-known to be dynamic even in response to images without dynamic content (Brincat & Connor, 2006)(Sugase, Yamane, Ueno, & Kawano, 1999)(Chen et al., 2014)(Meyer, Walker, Cho, & Olson, 2014), raising the question of what computations those neural activity dynamics might reflect. Prior work has proposed that such neuronal response dynamics could be the result of different types of circuits executing different types of computation such as: 1) recurrent circuits within each ventral stream processing stage implementing local normalization of the feedforward information as it passes through the stage (Carandini, Heeger, & Movshon, 1997)(Schwartz & Simoncelli, 2001)(Carandini & Heeger, 2012), 2) feedback circuits between each pair of ventral stream stages implementing the integration of top-down with bottom-up information to improve the current (online) inference (Seung, 1997)(Lee, Yang, Romero, & Mumford, 2002)(Zhang & Heydt, 2010)(Epshtein, Lifshitz, & Ullman, 2008), or 3) feedback circuits between each pair of stages comparing top-down and bottom-up information to compute prediction errors that guide changes in synaptic weights so that neurons are better tuned to features useful for future feedforward behavior (learning) (Rao & Ballard, 1999). Thus, neural dynamics may reflect the various adaptive computations (within-stage normalization, top-down Bayesian inference) or reflect the underlying error intermediates that could be generated during those processes (e.g. predictive coding).

These computationally motivated ideas can each be implemented as neural circuits to ask which idea best predicts response dynamics across the visual hierarchy. Here, our main goal was to look beyond the initial, feedforward response edge to see if we could disambiguate among dynamics that might result from stacked feedforward, lateral, and feedback operations. Rather than record from a single processing level, we measured the dynamics of neural signals across three hierarchical levels (pIT, cIT, aIT) within macaque IT. We focused on face processing subregions within each of these levels for three reasons. First, prior evidence argues that these three subregions are tightly anatomically and functionally connected and that the subregion in pIT is the dominant input to the higher subregions (Grimaldi, Saleem, & Tsao, 2016)(Moeller, Freiwald, & Tsao, 2008). Second, because prior work argues that a key behavioral function of these three subregions is to distinguish faces from non-faces, this allowed us to focus our testing on a relatively small number of images targeted to engage that processing function. Third, prior knowledge of pIT neural tuning properties (Issa & DiCarlo, 2012) allowed us to design images that were quantitatively matched in their ability to drive neurons in the pIT input subregion but that should ultimately be processed into two separate groups (face vs. non-face). We reasoned that these images would force important computations for disambiguation to occur somewhere between the pIT subregion and the higher level (cIT, aIT) subregions. With this setup, our aim was to observe the dynamics at all three levels of the hierarchy in response to that image processing challenge so that we might discover – or at least constrain -- which type of computation is at work.

Consistent with the idea that the overall system performs face vs. non-face discrimination (i.e. face detection), we found that in the highest face processing stage (aIT), neurons rapidly developed and maintained a response preference for faces over non-faces even though our images were designed to be challenging for frontal face detection. However, we found that many neurons in the early (pIT) and intermediate (cIT) processing levels of IT had face selectivity that rapidly but paradoxically *decreased*. That is, the responses of these neurons evolved to prefer images of non-faces over images of faces within 30 milliseconds of their feedforward response. We found that standard feedforward models that employ local recurrences such as adaptation, lateral inhibition, and normalization could not capture this stage-wise pattern of image selectivity despite our best attempts. However, we found that *decreasing* -- rather than increasing -- face preference in early and intermediate processing stages is a natural dynamical signature of previously suggested “error coding” models (Rao & Ballard, 1999) in which the neural spiking activity at each processing stage carries both an explicit representation of the variables of interest (e.g. is a face present?) and an explicit encoding of errors computed between each pair of stages in the hierarchy (e.g. a face was predicted, but a non-face was present leading to an error).

## RESULTS

We leveraged the hierarchically arranged face processing system in macaque ventral visual cortex to study the dynamics of neural processing across a hierarchy (Tsao, Freiwald, Tootell, & Livingstone, 2006)(Tsao, Moeller, & Freiwald, 2008) (**Figure 1A**). The serially arranged posterior, central, and anterior face-selective subregions of IT (pIT, cIT, and aIT) can be conceptualized as building increasing selectivity for faces culminating in aIT representations (Freiwald & Tsao, 2010)(Chang & Tsao, 2017). Using serial, single electrode recording, we sampled neural sites across the posterior to anterior extent of the IT hierarchy in the left hemispheres of two monkeys to generate neurophysiological maps (**Figure 1A**; example neurophysiological map in one monkey using a faces versus non-face objects screen set) (Issa, Papanastassiou, & DiCarlo, 2013). We localized the recording locations *in vivo* and co-registered across all penetrations using a stereo microfocal x-ray system (~400 micron *in vivo* resolution) (Cox, Papanastassiou, Oreper, Andken, & DiCarlo, 2008)(Issa, Papanastassiou, Andken, & DiCarlo, 2010) allowing accurate assignment of sites to different face processing stages (n = 633 out of 1891 total sites recorded were assigned as belonging to a face-selective subregion based on their spatial location; see **Methods**). Results are reported here for sites that were spatially located in a face-selective subregion, that showed visual drive to any category in the screen set (see **Methods**), and that were subsequently tested with our face versus non-face challenge set (**Figure 1B**, left panel) (n = 115 pIT, 70 cIT, and 40 aIT sites).

**Figure 1.**
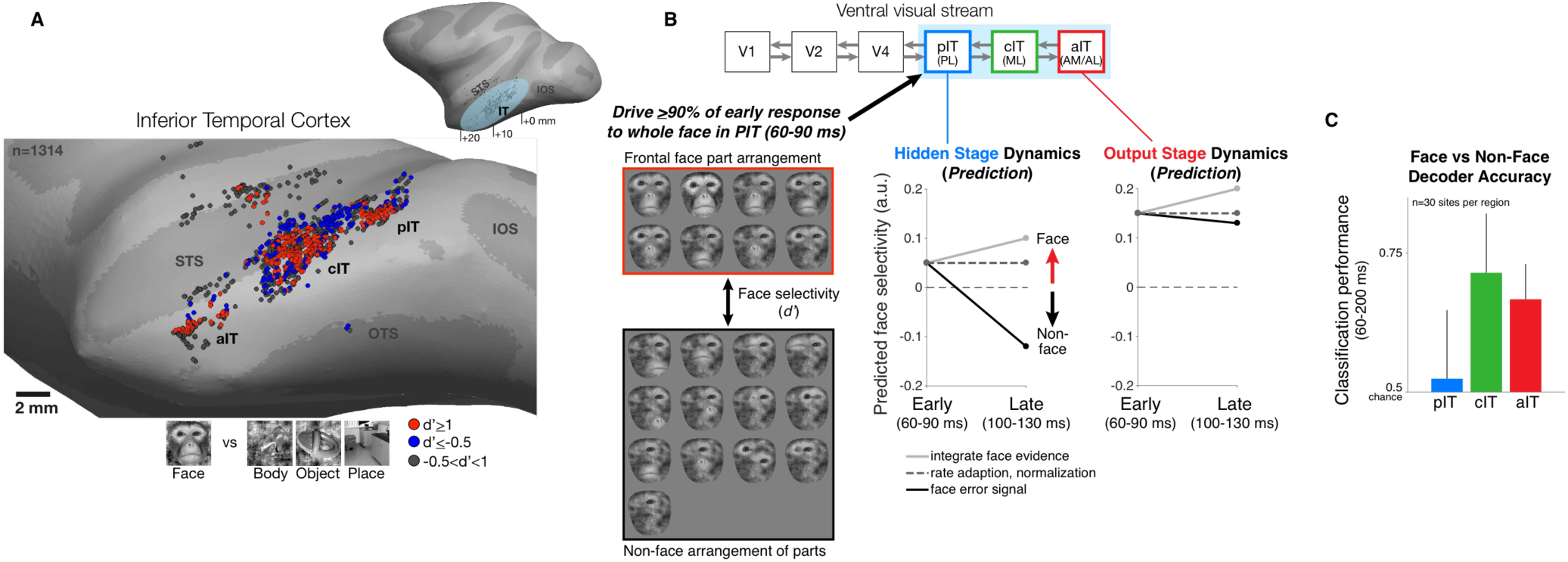
Neural recordings and experimental design in face-selective subregions of the ventral visual stream. **(A)** Neurons were recorded along the lateral convexity of the inferior temporal lobe spanning the posterior to anterior extent of IT (+0 to +20 mm AP, Horsely-Clarke coordinates) in two monkeys (data from monkey 1 are shown). Based on prior work, face-selective sites (red) were operationally defined as those with a response preference for images of frontal faces versus images of non-face objects (see **Methods**). While these neurons were found throughout IT, they tended to be found in clusters that mapped to previously identified subdivisions of IT (posterior, central, and anterior IT) and corresponded to face-selective areas identified under fMRI in the same subjects (Issa & DiCarlo, 2012)(Issa et al., 2013) (STS = superior temporal sulcus, IOS = inferior occipital sulcus, OTS = occipitotemporal sulcus). **(B)** (top diagram) The three visual processing stages in IT lie downstream of early visual areas V1, V2, and V4 in the ventral visual stream. (left) We designed our stimuli to focus on the intermediate stage pIT by seeking images of faces and images of non-faces that would, on average, drive equally strong initial responses in pIT. Novel images were generated from an exemplar monkey face by positioning the face parts in different positions within the face outline. This procedure generated both frontal face and non-face arrangements of the face parts, and we identified 21 images (red and black boxes) that drove the mean, early (60-100 ms) pIT population response to ≥90% of its response to the intact face (first image in red box is synthesized whole face; compare to the second image which is the original whole face), and of these 21 images, 13 images contained non-face arrangements of the face parts. For example, images with an eye centered in the outline (black box, 3^rd^ and 4^th^ rows) as opposed to the lateralized position of the eye in a frontal face (red box) have a global interpretation (“cyclops”) that is not consistent with a frontal face but still evoked strong pIT responses. Selectivity of neural sites (see **Figs. 3 & 4**) for face versus non-face images was quantified using a d’ measure. (middle) Computational hypotheses of cortical dynamics make differing predictions about how neural selectivity in pIT may evolve following an initial, weak preference signal for images of frontal faces. (right) Predictions of how aIT would behave as an output stage building selectivity for images of frontal faces through multiple stages of processing. **(C)** A population decoder, trained on average firing rates (60-200 ms post image onset, linear SVM classifier) for frontal face versus non-face arrangements of the face parts in this image subset, performed poorly in pIT on held-out trials of the same images (trial splits used so that the same images were shown in classifier training and testing). However, the particular class (face vs non-face) could be determined at above chance levels when reading the cIT and aIT population responses.

Our experimental design was intended to test previously proposed computational hypotheses of hierarchical neural dynamics during visual face processing (**Figure 1B**). Briefly, these hypotheses predict how stimulus preference (in this instance, for faces versus non-faces) might change over time in a neural population (**Figure 1B**, middle panel): (1) simple spike-rate adaptation predicts that initial rank-order selectivity (i.e. relative stimulus preference) will be largely preserved (**Figure 1B**, dashed line) while neurons adapt their absolute response strength over time, (2) local normalization predicts that stronger responses are in some cases normalized to match weaker responses based on population activity to specific dimensions (Carandini et al., 1997); importantly, normalization is strongest for nuisance (non-coding) dimensions (e.g. low versus high stimulus contrast) and in its idealized form would not alter selectivity along coding dimensions (e.g. face versus non-face) (**Figure 1B**, dashed line), (3) evidence accumulation through temporal integration, winner-take-all through recurrent inhibition, or Bayesian inference through top-down feedback mechanisms all predict increasing selectivity for faces over time (Lee & Mumford, 2003) (**Figure 1B**, light gray line), and (4) predictive coding posits that, for neurons that are coding error, their responses would show increasing activity for non-faces (images whose properties are inconsistent with predictions of a face) and thus decreasing face selectivity over time (Rao & Ballard, 1999) (**Figure 1B**, black line). Note, that error signaling is a qualitatively different computation than normalization, as error coding predicts a decreased response along the coding dimension (face versus non-face) whereas normalization would ideally not affect selectivity for faces versus non-faces and only affect variation along orthogonal, nuisance dimensions. Properly testing these predictions (no change in face selectivity, increased face selectivity, decreased selectivity to be anti-face preferring) requires measurements from the intermediate stages of the hierarchy as all of these models operate under the premise that the system builds and maintains a preference for faces at the top of the hierarchy (**Figure 1B**, right, and see Introduction). Thus, the intermediate stages (here pIT, see **Figure 1B**) are most likely to be susceptible to face/non-face confusion and thus be influenced by, for example, the top-down mechanisms posited in Bayesian inference and predictive coding where higher areas encode the face predictions that directly influence the responses of lower areas (Lee & Mumford, 2003)(Rao & Ballard, 1999).

### Face and non-face images driving similar initial responses in pIT

Here, we chose to focus our key, controlled tests on pIT – an intermediate stage in the ventral stream hierarchy, but the first stage within IT where neural specialization for face detection (i.e. face vs. non-face) has been reported (Grimaldi et al., 2016). Consistent with its intermediate position in the ventral visual system, we had previously found that pIT face-selective neurons are not truly selective for whole faces but respond to local face features, specifically those in the eye region (Issa & DiCarlo, 2012). Taking advantage of this prior result, we created both face and non-face stimuli that challenged the face processing system by strongly driving pIT responses, thus forcing the higher IT stages to complete the discrimination between face and challenging non-face images. To generate ambiguous face-like images, we systematically varied the positions of parts, in particular the eye, within the face (Issa & DiCarlo, 2012) (see **Methods**). This set included images that contained face parts in positions consistent with a frontal view of a face or images that only differed in the relative spatial configuration of the face parts within the face outline (**Figure 1B**, left). Of the 82 images screened, we identified 21 part configurations that each drove the pIT population response to ≥90% of its response to a correctly configured whole face. Of those 21 images, 13 images were inconsistent with the part configuration of a frontal face (**Figure 1B**, black box). For the majority of the results that follow, we focus on comparing the neural responses to these 13 pIT-matched images that could *not* have arisen from frontal faces (referred to hereafter as “non-face images”) with the 8 images that could have arisen from frontal faces (referred to hereafter as “face images”). Again, we stress that these two groups of images were selected to evoke closely matched initial pIT population activity.

Importantly, the pIT-matched images used here presented a more stringent test of face vs. non-face discrimination than prior work. Specifically, most prior work used images of faces and non-face objects (“classic images”) that contain differences across multiple dimensions including local contrast, spatial frequency, and types of features (Tsao et al., 2006)(Afraz, Kiani, & Esteky, 2006)(Moeller, Crapse, Chang, & Tsao, 2017)(Sadagopan, Zarco, & Freiwald, 2017). Consistent with this, we found that the population decoding classification accuracy of our recorded neural populations using these classic images (faces versus non-face objects) is near perfect (>99% in pIT, cIT, and aIT, n=30 sites per region). However, we found that population decoding classification accuracy for the pIT-matched face vs. non-face images we used here was near chance level (50%) in pIT (**Figure 1C**, blue bar; by comparison, classification accuracy for face versus non-face objects classification was 99.6% using the same pIT sites). Further downstream in regions cIT and aIT, we found that the linear population decoding classification of these pIT-matched face vs. non-face images was well above chance, suggesting that our pIT-matched face detection challenge is largely solved somewhere between pIT and aIT (**Figure 1C**).

### Time course of responses in pIT for images with face versus non-face arrangements of parts

We next closely examined the pIT neural response dynamics. To do this, we defined a face preference value (d’; see Methods) that measured each site’s average selectivity for the face images relative to the non-face images, and we asked how a given site’s preference evolved over time (see alternative hypotheses in **Figure 1B**). First, we present three example sites which were chosen based on having the largest selectivity (absolute d’) in the late phase (100-130 ms post image onset). In particular, most standard interpretations of face processing would predict a late phase preference for faces (d’ > 0). However, all three sites with the largest absolute d’ had evolved a strong late phase preference for the non-face images (d’ < 0) despite having had very similar rising edge responses to the face and non-face stimulus classes (response in early phase from 60-90 ms) (**Figure 2**, left column). A late, non-face preference was not restricted to the example sites as a majority of pIT sites (66%) preferred non-faces over faces in the late response phase (prefer frontal face arrangement: 60-90 ms = 66% vs. 100-130 ms = 34%; p = 0.000, n = 115) (**Figure 3B**, blue bars).

**Figure 2.**
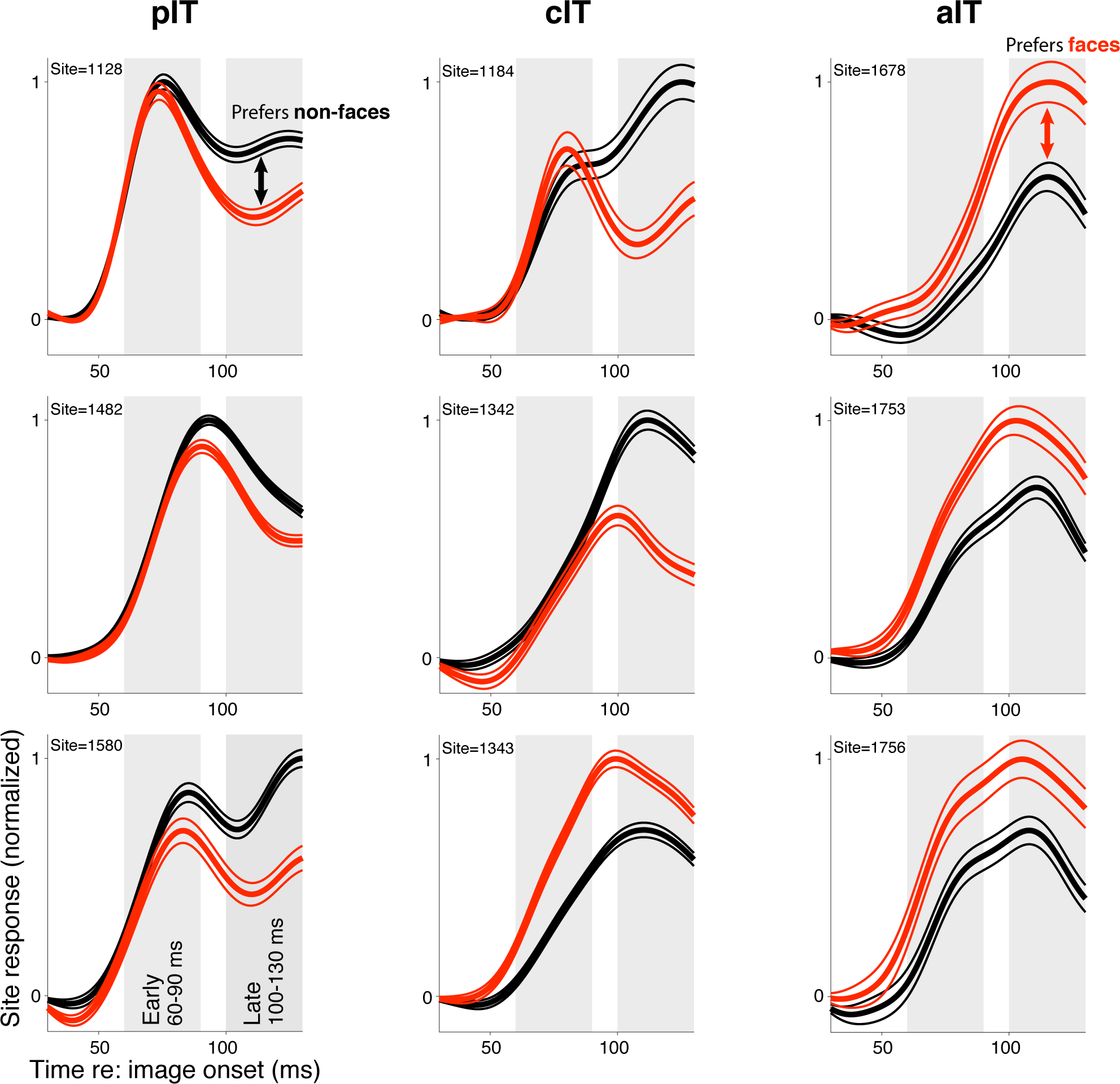
Responses in example sites to face-like images with typical and atypical face part arrangements. The three sites with the highest selectivity in the late response phase in each region are shown (pIT, cIT, and aIT; left, middle, and right columns, respectively) (d’ selectivity measured in a 100-130 ms window, gray shaded region shown in bottom, left panel). While the three aIT sites (right column) demonstrated late phase selectivity for face images, the three pIT sites evolved the opposite preference in their late phase (100-130 ms) responses (red line = mean response of 8 images shown in **Figure 1B** red box, and black line = mean response of 13 images shown in **Figure 1B** black box).

**Figure 3.**
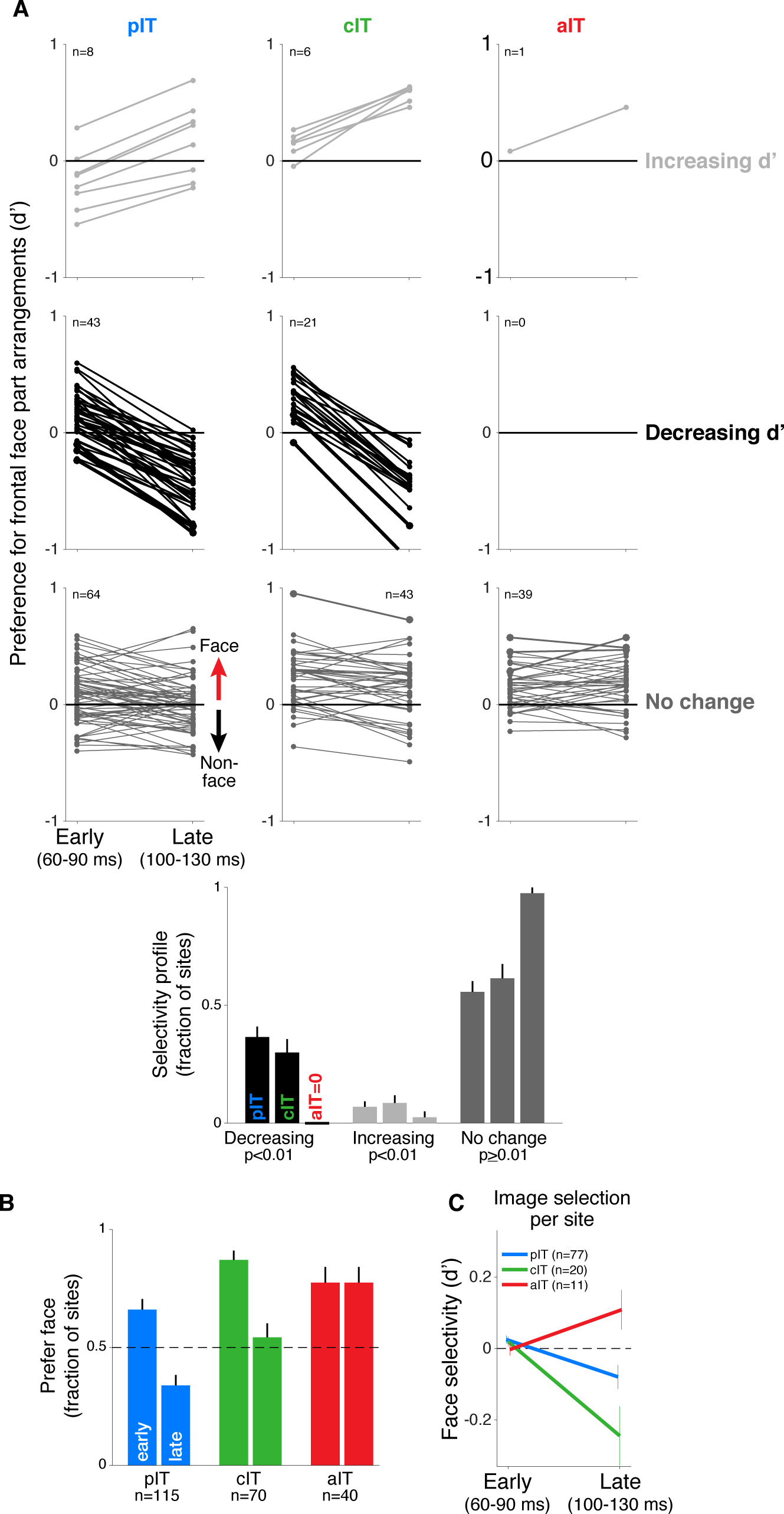
Time course of neural response preferences in pIT, cIT, and aIT for images with face versus non-face arrangements of the parts. **(A)** Preferences for frontal face versus non-face part arrangements for each site are plotted in both early (60-90 ms post image onset) and late (100-130 ms) time windows. Sites are grouped based on region (pIT, cIT, aIT) and whether they showed a significant change in selectivity from early to late time windows (light gray = increased preference, black = decreased preference, and dark gray = no change in preference for face versus non-face images, significance tested at p < 0.01 level; example sites from **Figure 2** are plotted using thicker, darker lines). Many sites in pIT and cIT showed a decreasing preference for frontal face versus non-face images over time (black lines, middle row, left and center panels). In contrast, no sites in aIT had this dynamic (middle row, right panel). **(B)** The fraction of sites whose responses showed a preference for images of typical, face-like arrangements of the face parts in pIT (blue), cIT (green) and aIT (red) in the early (60-90 ms) and late (100-130 ms) phase of the response. Note that, in the late phase of the response, most pIT neurons paradoxically showed a preference for non-face arrangements of face parts. **(C)** Selectivity measured for images driving similar responses within a site. This procedure ensured matched initial responses on a site-by-site basis rather than using a fixed set of images based on the overall population response (i.e. the fixed image set of **Figure 1B**; here, the initial d’ for 60-90 ms is close to zero when images are selected site by site). Although initial response differences were near zero when using site based image selection, a late phase preference for non-face images still emerged in pIT and cIT but not in aIT similar to the decreasing selectivity profile observed when using a fixed image set for all sites.

Next, we examined the dynamics of face selectivity across the pIT population as this is key to disambiguating among the competing models outlined earlier (**Figure 1B**). In the adaptation and normalization models, we would expect no change in the average population face selectivity, and the evidence accumulation, winner-take-all, or Bayesian inference models predict an increase in face selectivity over the population over time. Instead, we found that many sites significantly decreased their face preference over time similar to the three example sites. Of the 51 sites in our pIT sample that showed a significantly changing preference over time (p < 0.01 criterion for significant change in d’), 84% of these sites showed a decreasing preference (n = 43 of 51 sites, p < 10^-6, binomial test, n = 115 total sites) (**Figure 3A**, left column, light gray vs. black lines). This surprising trend -- decreasing face preference -- was strong enough that it erased any small, initial preference for images of frontally arranged face parts over the population (median d’: 60-90 ms = 0.11 ± 0.02 vs. 100-130 ms = -0.12 ± 0.03, p = 0.000, n=115 sites), and this trend was observed in both monkeys when analyzed separately (p_M1_ = 0.000, p_M2_ = 0.002, n_M1_ = 43, n_M2_ = 72 sites; **Figure 4A**). This decreasing face selectivity over time was driven by decreasing firing rates to the face images containing normally arranged face parts. Responses to these images were weaker by 18% on average in the late phase of the response compared to the early phase (Δrate (60-90 vs 100-130 ms) = -18% ± 4%, p = 0.000; n = 7 images) while responses to the non-face images with atypical spatial arrangements of face parts -- also capable of driving high early phase responses -- did not experience any firing rate reduction in the late phase of the response (Δrate (60-90 vs 100-130 ms) = 2 ± 1%, p = 0.467; n = 13 images).

**Figure 4.**
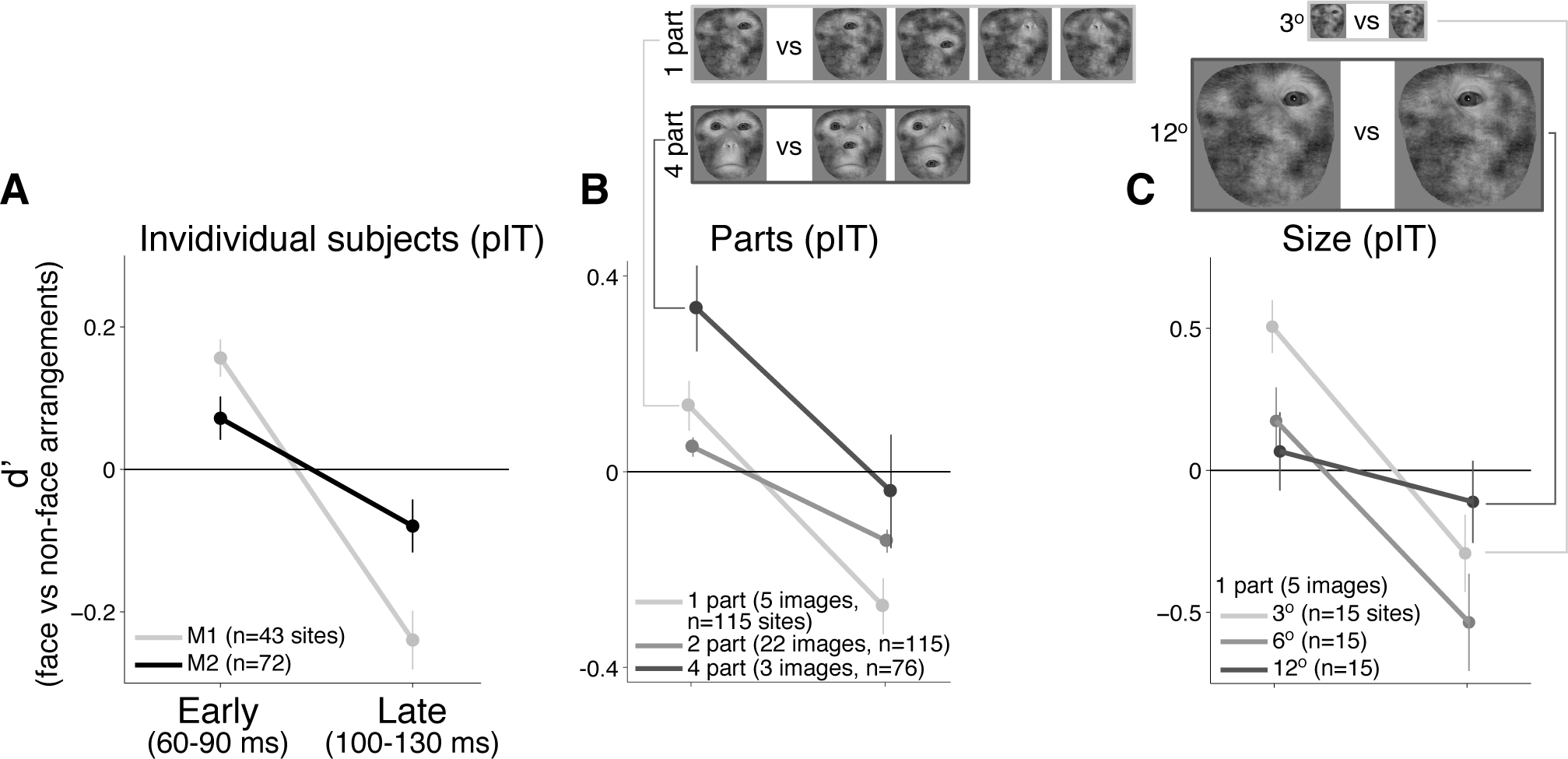
Individual monkey comparison and image controls for the decreasing selectivity profile in pIT. **(A)** Preference for images with frontal face part arrangements analyzed separately for each monkey. Median d’ of pIT sites in both early and late time windows is shown. **(B)** Preference for images with face versus non-face arrangements of the parts was re-computed using image subsets containing the same number of parts in the outline (the five 1-part and the three 4-part image subsets shown at top; the larger 2-part subset contained 30 images and is not shown). **(C)** The 1-part image subset was further tested at three different sizes (3°, 6°, and 12°). In all cases, pIT responses showed a decreasing preference over time for typically-arranged face parts leading to a preference for atypically arranged face parts in the later time window (100-130 ms).

The above observation of decreasing face preference over the pIT population seemed most consistent with the prediction of error coding models, but one potential confound was that initial responses to faces and challenging non-faces were not perfectly matched across the population (recall that we only required face and non-face images to drive a response ≥90% of the whole face response). As a result, initial selectivity was non-zero (d’ = 0.11, n=115 sites). This residual face preference may be small, but if this residual face selectivity is driven by nuisance dimensions, for example excess stimulus contrast in the face class relative to the non-face class, then the face class may have experienced stronger activity dependent adaptation or normalization resulting in a decreasing face preference over time. To more adequately limit general activity dependent mechanisms leading to decreasing face responses, we performed control analyses where initial activity was tightly matched per site or where the number of parts were matched across images.

### Controls in pIT for firing rate and low-level image variation

To strictly control for the possibility that simple initial firing rate differences could predict the observed phenomenon, we re-computed selectivity after first matching initial responses site-by-site. For this analysis, images were selected on a per site basis to evoke similar initial firing rates (images driving initial response within 25% of synthetic whole face response for that site, at least 5 images required per class). This image selection procedure virtually eliminated any differences in initial responses between the face and non-face image classes and hence any firing rate difference driven by differences in nuisance parameters between faces and challenge non-faces (**Figure 3C**, 60-90 ms), yet we still observed a significant drop in preference for images with typical frontal face part arrangements versus atypical face part arrangements in pIT (Δ_d’_ = -0.10 ± 0.03, p = 0.001, n = 77) (**Figure 3C**, blue line). Thus, the remaining dependence of firing rate dynamics on the image class and not on initial response strength argued against an exclusively activity based explanation to account for decreasing neural responses to faces over time. Further arguing against this hypothesis, we found that the pattern of late phase population firing rates in pIT across images could not be significantly predicted from early phase pIT firing rates for each image (ρ_pIT early, pIT late_ = 0.07 ± 0.17, p = 0.347; n = 20 images).

Thus far, we have performed analyses where images from the face and non-face class were similar in their initially evoked response which equated images at the level of neural activity but produced images varying in the number of parts. An alternative is to match the number of face parts between the face and non-face classes as another means of limiting the differences in nuisance dimensions such as the contrast, spatial frequency and retinal position of energy across images (see examples in **Figure 4B**). When we recomputed selectivity across subsets of images containing a matched number of one, two, or four parts (n=5, 30, and 3 images, respectively), we still observed that pIT face selectivity decreased. For all three image subsets controlling the number of face parts, d’ began positive on average in the sampled pIT population (i.e. preferred frontal face part arrangements in 60-90 ms post-image onset) (median d’ for 60-90 ms = 0.13 ± 0.05, 0.05 ± 0.02, 0.33 ± 0.09 for one, two, and four parts) and significantly decreased in the next phase of the response (100-130 ms post-image onset) becoming negative on average (median d’ for 100-130 ms: -0.27 ± 0.06, -0.14 ± 0.02, -0.04 ± 0.12; one, two, four parts: p = 0.000, 0.000, 0.004, for d’ comparisons between 60-90 ms and 100-130 ms, n = 115, 115, 76 sites) (**Figure 4B**). A similar decreasing face selectivity profile was observed when we re-tested single part images at smaller (3°) and larger (12°) image sizes suggesting a dependence on the relative configuration of the parts and not on their absolute retinal location or absolute retinal size (median d’ for 60-90 ms vs. 100-130 ms: three degrees = 0.51 ± 0.09 vs. -0.29 ± 0.14, twelve degrees = 0.07 ± 0.14 vs. -0.11 ± 0.14; n = 15; p = 0.000, 0.025, 0.07) (**Figure 4C**). Thus, we suggest that the decreasing population face selectivity dynamic in pIT is a fairly robust phenomenon specific to the face versus non-face dimension as this dynamic persists even when limiting potential variation across nuisance dimensions. This phenomenon can be distinguished from normalization mechanisms that would operate most strongly to reduce selectivity along nuisance dimensions rather than along dimensions that directly solve the face versus non-face discrimination challenge.

### Time course of responses in aIT and cIT for images with face versus non-face arrangements of parts

Under the possibility that decreasing face selectivity in intermediate stage pIT is a signature of error signaling, we next asked whether the source of the prediction signal could be observed in higher cortical areas. In the anterior face-selective regions of IT which are furthest downstream of pIT and reflect additional stages of feedforward processing (see block diagram in **Figure 1B**), we did not observe the strong decreases in the selectivity profile seen in pIT. Indeed, the three sites with the greatest selectivity (absolute d’) in the late response phase (100-130 ms) in our aIT sample all displayed a preference for frontal face part arrangements (d’ > 0) (**Figure 2**, right column). Also, in contrast to the dynamic selectivity profiles observed in many pIT sites, 98% of aIT sites (39 of 40) did not significantly change their relative preference for face vs. non-face arrangements of the parts (p < 0.01 criterion for significant change at the site level) (**Figure 3A**, right column, bottom row, dark gray sites). Rather, we observed a stable selectivity profile over time in aIT (median d’: 60-90 ms = 0.13 ± 0.03 vs. 100-130 ms = 0.17 ± 0.03, p = 0.34, n=40 sites). As a result, the majority of anterior sites preferred images with typical frontal arrangements of the face parts in the late phase of the response (prefer face: 60-90 ms = 78% of sites vs. 100-130 ms = 78% of sites; p = 0.451, n = 40 sites; **Figure 3B**, red bars) despite only a minority (34%) of upstream sites in pIT preferring these images in their late response. Thus, spiking responses of individual aIT sites were as expected from a computational system whose purpose is to detect faces, as previously suggested (Freiwald & Tsao, 2010), and aIT serves as one candidate for the putative prediction signal underlying face prediction errors.

In cIT whose anatomical location is intermediate to pIT and aIT, we observed many sites with decreasing selectivity (**Figure 2C & 3A**, middle columns), a dynamic that persisted even when we tightly matched initial responses on a site by site basis similar to pIT (**Figure 2C**, green line). The overall stimulus preference in cIT was intermediate to that of pIT and aIT (**Figure 3B**) consistent with the intermediate position of cIT in the IT hierarchy. Interestingly, we found that the patterns of responses across images in the early phases of cIT and aIT activity were significant predictors of late phase activity in pIT (ρ_cIT early, pIT late_ = -0.52 ± 0.11, p = 0.000; ρ_aIT early, pIT late_ = -0.36 ± 0.14, p = 0.012; n_pIT_=115, n_cIT_=70, n_aIT_=40 sites; n = 20 images), even better predictors than early phase activity in pIT itself (ρ_pIT early, pIT late_ = 0.07 ± 0.17, p = 0.347). That is, for images that produced high early phase responses in cIT and aIT, the following later phase responses of units in the lower level area (pIT) tended to be low, consistent with error coding models which posit that feedback from higher areas (in the form of predictions) would contribute to the decreasing selectivity observed in lower areas.

### Computational models of neural dynamics in IT

We next proceeded to formalize the conceptual ideas introduced in **Figure 1B** and build neurally mechanistic, dynamical models of gradually increasing complexity to determine the minimal set of assumptions that could capture our empirical findings of non-trivial, dynamic selectivity changes during face detection across face-selective subregions in IT. Previous functional and anatomical data show that the face-selective subregions in IT are connected forming an anterior to posterior hierarchy and show that pIT serves as the primary input into this hierarchy (Moeller et al., 2008)(Freiwald & Tsao, 2010)(Grimaldi et al., 2016). Thus, we evaluated dynamics in different hierarchical architectures using a linear dynamical systems modeling framework where pIT, cIT, and aIT act as sequential stages of processing (**Figure 5** and see **Methods**). A core principle of feedforward ventral stream models is that object selectivity is built by stage-wise feature integration in a manner that leads to relatively low dimensional representations at the top of the hierarchy abstracted from the high-dimensional input layer. We were interested in how signals temporally evolve across a similar architectural layout. We used the simplest feature integration architecture where a unit in a downstream area linearly sums the input from units in an upstream area, and we stacked this computation to form three layer networks (**Figure 5B**). This simple, generic feedforward encoding model conceptualizes the idea that different types of evidence, local and global (i.e. information about the parts and the relative spatial arrangement of parts), have to converge and be integrated to separate face from non-face images in our image set. We used linear networks as monotonic nonlinearities can be readily accommodated in our framework (Seung, 1997)(Rao & Ballard, 1999)(also see **Figure 7C**). Importantly, we used a simple encoding scheme as our goal was not to build full-scale deep neural network encoding models of image representation (Yamins et al., 2014) but to bring focus to an important biological property that is often not considered in deep nets, neural dynamics.

**Figure 5.**
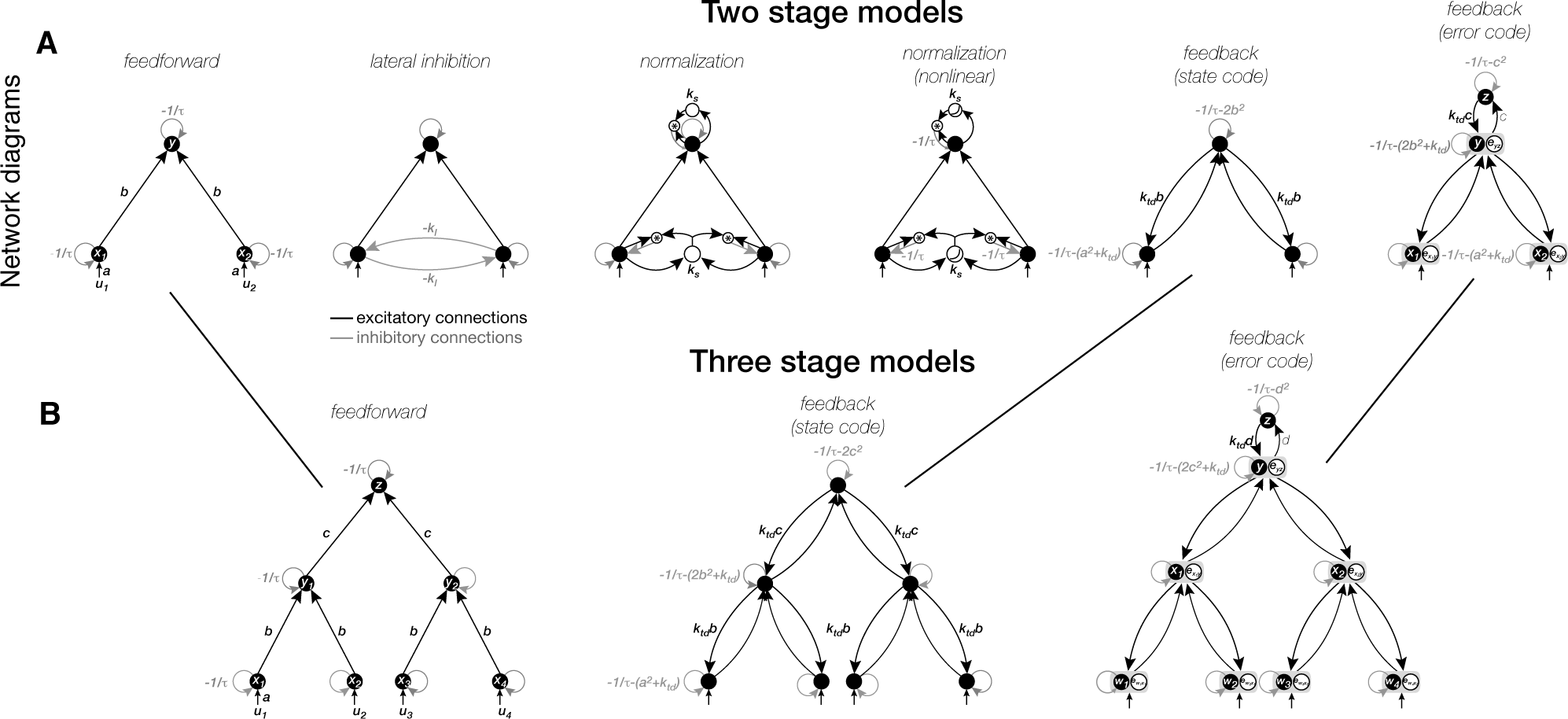
Model diagrams. **(A)** Network diagrams of two-stage models used in **Figure 7A,B**. The base feedforward architecture and parameters are shown in the first column while recurrent models with added connections and parameters are shown in the remaining columns. All models have two inputs (*u*_*1*_, *u*_*2*_), at least two hidden units (*x*_*1*_, *x*_*2*_) (a.k.a. layer 1 of the network), and at least one output unit (*y*). For simulations, the inputs *u*_*1*_, *u*_*2*_ are set independently to simulate each hidden node receiving different amounts of external drive, depending on the choice of the applied image relative to the unit’s preferred image. The connection weights B = [*b*_*1*_, *b*_*2*_] transforming the hidden stage activations to the output unit are modeled as the same (*b*_*1*_ = *b*_*2*_). All units have self-connections that determine the degree of leak current set by the time constant *τ*. In the normalization models, the leak term is additionally controlled (linearly or nonlinearly) by the total activity in each stage (third and fourth columns). In the feedback-based model, the feedback connections are symmetric to the feedforward connections with weights *B*^*T*^ = [*b*_*1*_, *b*_*2*_]^*T*^, a column vector (fifth and sixth columns). The error coding feedback model (sixth column) has an additional stage that contributes to computation of error in the second stage (see **Methods** for details). **(B)** Extensions of the two-stage model architectures to three stages are shown only for the feedforward and feedback models (compare to two-stage diagrams in **(A)**). The three-stage model has an additional hidden processing stage compared to the two-stage model. An extra node is introduced at the top of the hierarchy to produce an error signal in the third stage of the error coding model.

**Figure 7.**
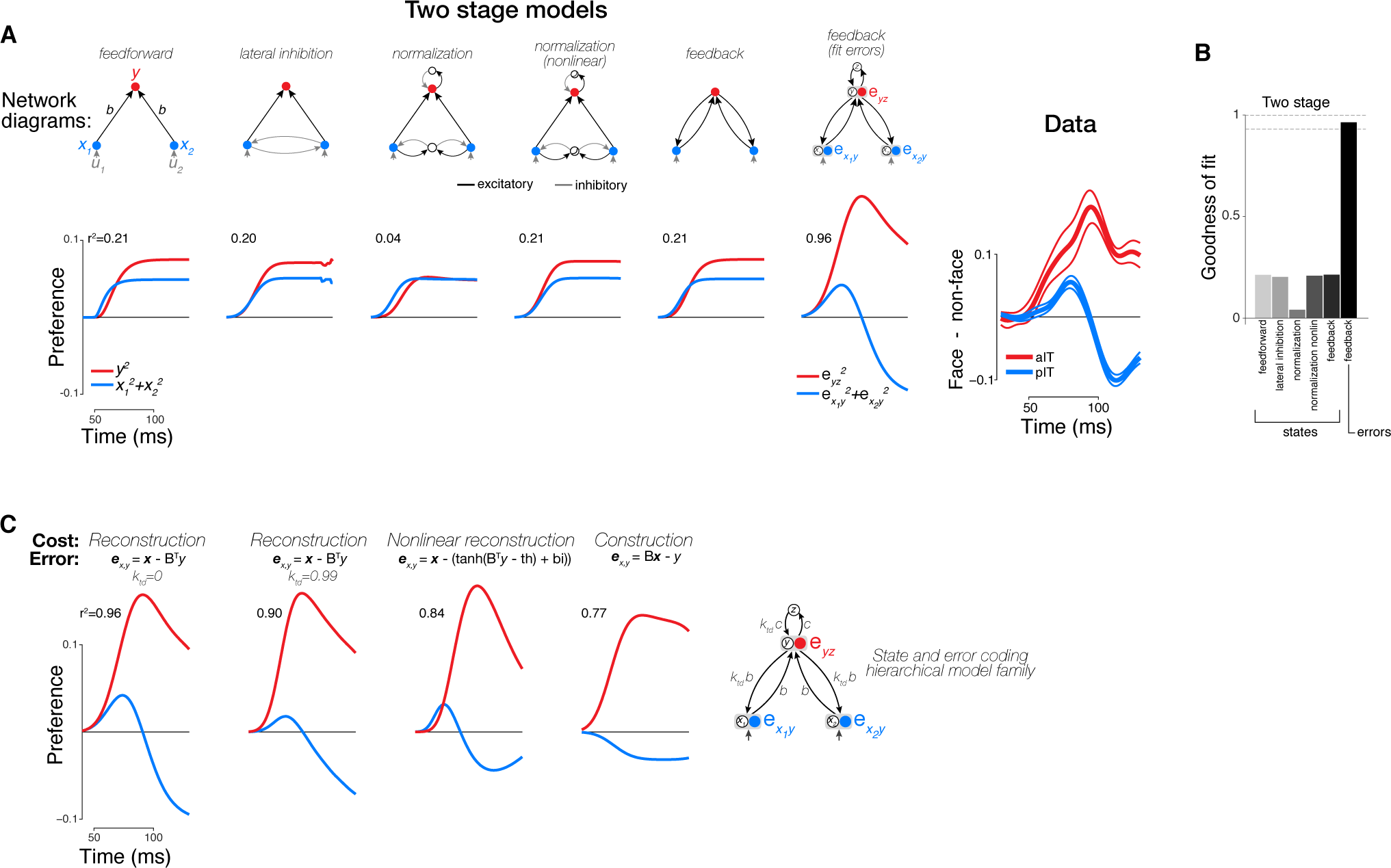
Two stage model fits to neural dynamics and comparison of variants of error coding hierarchical models that use different algorithms for online inference. **(A)** We tested how well two stage models could reproduce decreasing selectivity observed in the hidden layer (same format as **Figure 6A**). The two stage error coding feedback model displayed similar dynamical phenomena as the data suggesting that three stages of processing were not necessary to obtain error signal dynamics. **(B)** Goodness of fit to population neural data (right plot in **(A)**) for all two stage models. **(C)** Existing forms of error computing networks can be distinguished by the type of online inference algorithm that they use. In one case, inference does not utilize top-down information between stages (classic error backpropagation; between-stage feedback connections shown are not used in these networks during runtime). On the other hand, between-stage feedback can be used such as in more general forms of error backpropagation and predictive coding. We approximated these two extremes by including a parameter (*k*_*td*_, see **Methods**) controlling the relative weighting of bottom-up (feedforward) and top-down (feedback) evidence during online inference (first and second panels). We found that top-down inference between stages was not necessary to produce the appropriate error signal dynamics, and *k*_*td*_ was equal to zero in our best fitting two-layer and three-layers models (first panel is same model as two-layer error coding model in **(A)**) although models with *k*_*td*_ ~ 1 also performed well (second panel). Models can also differ in their goal (cost function) which directly impacts the error signals required (equations in top row). Under a nonlinear reconstruction goal (emulating the nonlinear nature of spiking output), the resulting error signals were still consistent with our data (third column). A simple sigmoidal nonlinearity, however, did lead to additional details present in our neural data such as a rapid return of stimulus preference to zero in the hidden layer. When we tested a discriminative, construction goal more consistent with a supervised learning setting (e.g. classic error backpropagation) where bottom-up responses simply have to match a downstream target signal in classification tasks, we found that the errors of construction did not match the data as well as reconstruction errors (compare fourth column to first three columns) although both types of error outperformed all state-based models and were overall very similar (compare to **(A)**).

We implemented a range of ideas previously proposed in the literature. The functional implications of these ideas were highlighted in **Figure 1B**, but at a mechanistic level, these functional properties can be directly realized via different recurrent processing motifs between neurons (**Figure 5**, base feedforward model (first column) is augmented with recurrent connections to form new models (remaining columns)). For example, self-connections can be viewed as implementing spike rate adaptation in a feedforward architecture (**Figure 5A**, top row), lateral connections support winner-take-all or normalization mechanisms (Carandini et al., 1997) (**Figure 5A**, second, third, and fourth columns), and top-down connections implement Bayesian inference (Seung, 1997) (**Figure 5A**, fifth and sixth columns). Here, normalization is implemented by a leak term that scales adaptation (the degree of decay in the response) and is controlled recursively by the summed activity of the network (Carandini et al., 1997), ultimately scaling down responses to strong driving stimuli over time. To constrain our choice of a feedback-based model, we took a normative approach minimizing a quadratic reconstruction cost between stages as the classical reconstruction cost is at the core of an array of hierarchical generative models including hierarchical Bayesian inference (Lee & Mumford, 2003), Boltzmann machines (Ackley, Hinton, & Sejnowski, 1985), analysis-by-synthesis networks (Seung, 1997), sparse coding (Olshausen & Field, 1996), predictive coding (Rao & Ballard, 1999), and autoencoders in general (Rifai, Vincent, Muller, Glorot, & Bengio, 2011). Optimizing a quadratic loss results in feedforward and feedback connections that are symmetric -- reducing the number of free parameters -- such that inference on the represented variables at any intermediate stage is influenced by both bottom-up sensory evidence and current top-down interpretations. Critically, a common feature of this large model family is the computation of between-stage error signals via feedback, which is distinct from state-estimating model classes (i.e. feedforward models) that do not compute or propagate errors. A dynamical implementation of such a network uses leaky integration of error signals which, as shared computational intermediates, guide gradient descent of the values of the represented variables to a previously learned target value (∆activity of each neuron => online inference) or descend the connection weights to values that give the best future behavior (∆synaptic strengths => offline learning), here defined as an unsupervised reconstruction goal (similar results were found using other goals and networks such as supervised discriminative networks; see **Figure 7C**).

When we fit each of the models to our neural data, they generally produced an increase in selectivity from the first stage of the network to the later stages of the network. This increase is not surprising because the models had built-in converging feedforward connections from the first to second to third stages (**Figure 6A**, first five columns, compare blue to red curves). However, discrepancies between models emerged in the lower stages where we found that neither the lateral inhibition model, nor the normalization model, could capture the decreasing selectivity phenomenon observed in pIT. Instead, the selectivity of these models simply increased to a saturation level set by the leak term (shunting inhibition) in the system (**Figure 6A**, first five columns). Similar behavior was present when we tried a nonlinear implementation of the normalization model that more powerfully modulated shunting inhibition (Carandini et al., 1997). That normalization proved insufficient to generate the observed neural dynamics can be explained by the fact that the normalized response to a stimulus cannot easily fall below the response to a stimulus that was initially similar in strength. Thus, a decreasing average preference for a stimulus across a population of cells (i.e. **Figures 2-4**, pIT data) for similar levels of average input is difficult when only using a basic normalization model mediated by surround (within-stage) suppression.

**Figure 6.**
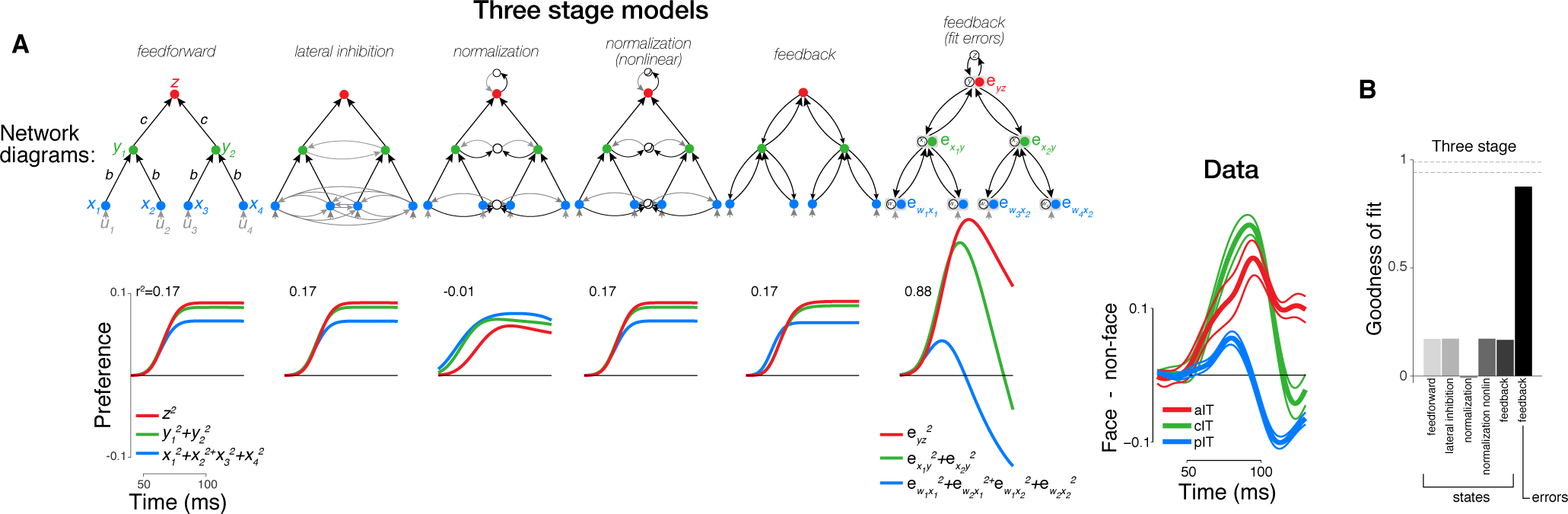
Computational modeling of neural dynamics in IT. **(A)** Three stage neural networks with recurrent dynamics (network diagrams in top row; see **Methods** and **Figure 5B**) were constructed to model neural signals measured in pIT, cIT, and aIT corresponding to the first (blue), second (green), and third (red) model processing stages. All models received four inputs (gray) into four hidden stage units (blue) which sent feedforward projections that converged onto two units in the next layer (green). Besides this feedforward architecture, additional excitatory and inhibitory connections between units were used to implement recurrent dynamics (self-connections reflecting leak currents are not shown here for clarity; see **Figure 5B** for detailed diagrams). In the five models on the left, the responses of the simulated neurons are assumed to code the current estimates of some set of features in the world (a.k.a states), as is standard in most such networks. The best fit to the population averaged neural data (far right) of the states of each model class are shown (first five columns). These state coding models generally showed increasing selectivity over time from hidden to output layers and did not demonstrate the strong decrease of stimulus preference in their hidden processing stage as observed in the pIT and cIT neural population (blue and green lines). However, the neurons coding errors in a feedback-based hierarchical model did show a strong decrease of stimulus preference in the hidden processing stage (sixth column; reconstruction errors instead of the states were fit directly to the data). This model which codes the error signals (filled circles) also codes the states (open circles) (network diagram in sixth column of top row). **Far right**, population averaged neural selectivity profile for difference between frontal face versus non-face arrangements (normalized by the mean population response to the whole face) used in model fitting. **(B)** Goodness of fit of all three stage models tested to population averaged selectivity profiles (dashed lines represent mean and standard error of reliability of neural data as estimated by bootstrap resampling).

In contrast to the above models, we found that the feedback model capable of computing hierarchical error signals naturally displayed a strong decrease of selectivity in a sub-component of its first processing stage -- qualitatively similar behavior to the selectivity decrease that we observed in many pIT and cIT neural sites. Specifically, this model displayed these dynamics in the magnitude of its reconstruction error signals but not in its state signals (the feature values) (**Figure 6A**, compare fifth and sixth columns). These error signals integrate converging state signals from two stages -- one above (prediction) and one below (sensory evidence). The term “error” is thus meaningful in the hidden processing stages where state signals from two stages can converge. The top nodes of a hierarchy receive little descending input and hence do not carry additional errors with respect to the desired computation; rather, top nodes convey the face predictions that influence errors in bottom nodes. This behavior in the higher processing stages is consistent with our observation of explicit representation of faces in aIT in all phases of the response (**Figures 2-3**) and with similar observations of decodable identity signals by others in all phases of aIT responses for faces (Meyers, Borzello, Freiwald, & Tsao, 2015) and objects (Hung et al., 2005)(Majaj et al., 2015). We also found similar error dynamics when using a simpler two-layer network as opposed to three layers suggesting that these error signal dynamics along with prediction signals emerge even in the simplest cascaded architecture (**Figure 7A**).

### Predictions of an error coding hierarchical model

While our neural observations at multiple stages of the IT hierarchy led us to the error coding hierarchical model above, a stronger test of the idea of error signaling is whether it predicts other IT neural phenomena. To identify stimulus regimes that would lead to insightful predictions, we asked in what way would the behavior of error-estimating hierarchical models differ most from the behavior of generic feedforward state-estimating models. Because our feedback-based model uses feedforward inference at its core, it behaves similarly to a state-estimating hierarchical feedforward model when the statistics of inputs match the learned feedforward weight pattern of the network (i.e. ‘natural’ images drawn from everyday objects and scenes) since for these inputs, feedforward inferences derived from the sensory data are aligned with top-down expectations. Thus, predictions of feedback-based models that could distinguish them from feedforward-only models are produced when the natural statistics of images are altered so that they differ from the feedforward patterns previously learned by the network and hence differ from the predictions generated in the network. We have (above) considered one such type of alteration: images where local face features are present but altered from their naturally occurring (i.e. statistically most likely) arrangement in frontal faces. Next, we tested two other image manipulations from recent physiology studies which yielded novel neural phenomena that lacked a principled, model-based explanation (Freiwald, Tsao, & Livingstone, 2009)(Meyer et al., 2014). To test whether the error coding hierarchical model family displays these behaviors, we fixed the architectural parameters derived from our fitting procedure in **Figure 6** and simply varied the input to this network, specifically the correlation between the inputs and the network’s feedforward weight pattern, in order to match the nature of the image manipulations performed in prior experiments.

#### Sublinear integration of the face features

In the face-selective subregion of cIT, the sum of the responses to the face parts exceeds the response to the whole face when averaging firing rates over a 200 ms window (Freiwald et al., 2009; see their Figure 2C), and we observed a similar phenomenon in the face-selective subregion in pIT (Issa & DiCarlo, 2012). Interestingly, examining the dynamics of part integration more closely within the first 100ms of the response reveals that the response to the whole face does begin at a level similar to the sum of the responses to the face parts but becomes sublinear in the late phase (whole face response relatively low compared to linear prediction from parts responses) (ratio of sum of responses to parts vs. response to whole: 60-90 ms = 1.5 ± 0.1, 100-130 ms = 4.6 ± 0.3; p = 0.000, n = 33 sites) (**Figure 8A**, left panel). This result runs counter to what would be expected in a model where selectivity for the whole face is built from the conjunction of the parts. In such a model, the population response to the whole face would be at least as large if not greater (superlinear) than the summed responses to the individual features. To test whether an error coding model exhibited the phenomenon of sublinear feature integration at the population level, we compared the response with all inputs active (co-occurring features) to the sum of the responses when each input was activated independently (individual features). The reconstruction errors in our feedback-based model showed a strong degree of sublinear integration of the inputs such that the response to the simultaneous inputs (whole) was much smaller than what would be predicted by a linear sum of the responses to each input alone (parts), and the model’s sublinear integration behavior qualitatively replicated the time course observed in pIT without any additional fitting of parameters (**Figure 8A**, right panel). Although we certainly expect that similar sublinear integration may also be observed in a normalization model, this would require a particularly strong form of normalization since population activity to the whole face would have to be normalized to nearly the same level as that for an individual part in the late response phase (ratio of response to single part vs. response to whole = 0.92 ± 0.06, 100-130 ms, n = 33 sites) despite being three times larger during the early response phase (ratio of response to single part vs. response to whole = 0.3 ± 0.02, 60-90 ms, n = 33 sites). Furthermore, the stacked normalization model could not account for the main dynamical phenomenon that we observed (**Figure 6**); therefore, an error coding perspective may provide a more parsimonious account of the observed set of dynamical phenomena.

**Figure 8.**
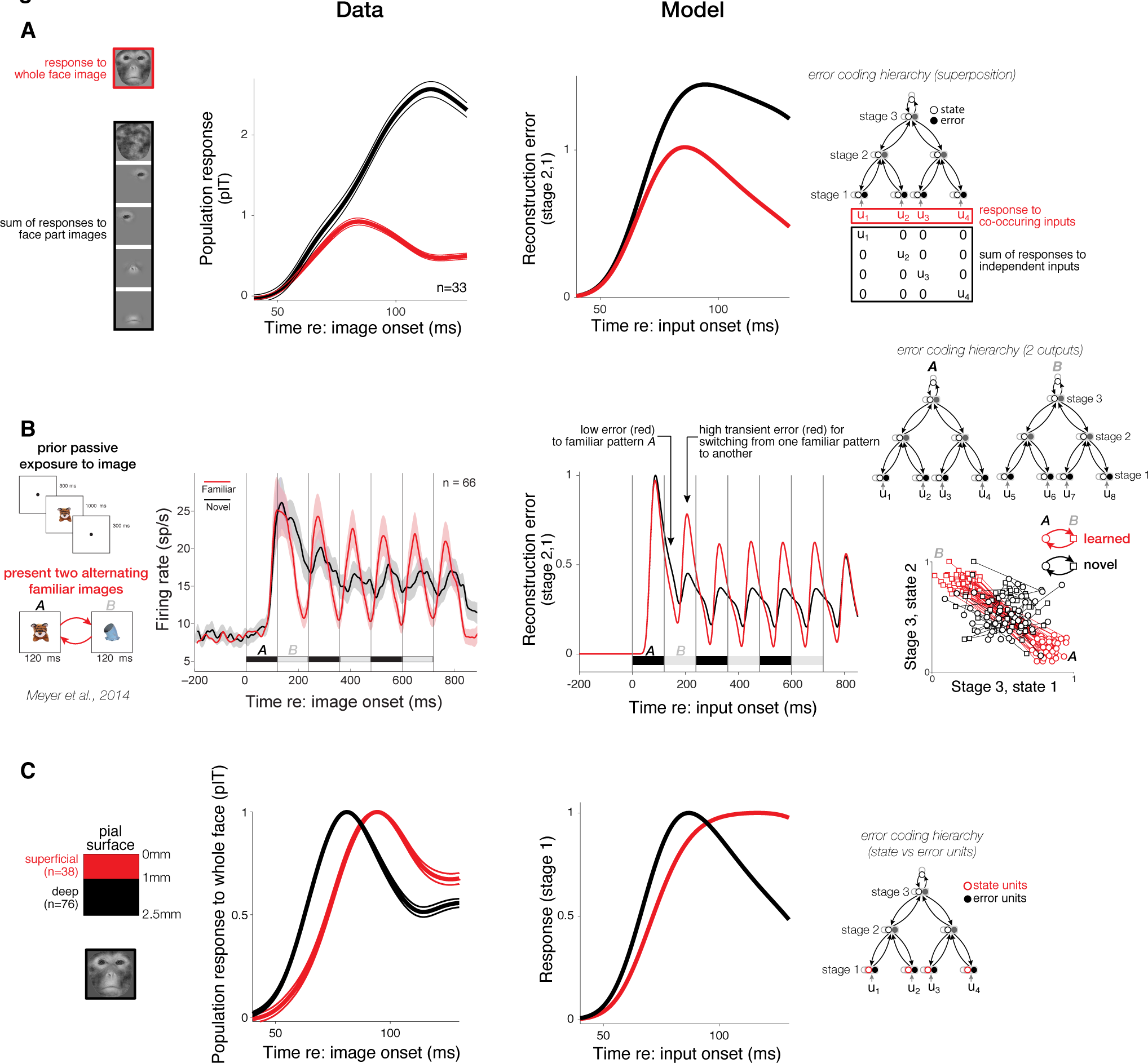
Comparison of neuronal phenomena to predictions of a feedback-based error coding model. To generate model predictions, the architectural parameters of the model (i.e. connection weights and time constants) were held fixed, and only the input patterns ***u*** were varied. **(A)** Neuronal (left panel): in pIT, we found that the sum of the neuronal response to the face parts presented individually (black) exceeded the response to the same parts presented simultaneously (i.e. a whole face, red). Each line is the mean response of 33 pIT sites normalized by the peak response to the whole face. Model (right panel): The magnitude of errors between stage 1 and stage 2 of the model showed a similar degree of sublinear integration (responses were normalized by peak response to the simultaneous input condition). **(B)** We extended the model to include two units in the third, output stage that responded to two learned input patterns (see **Methods**) with increased separation of patterns *A* and *B* in this high-level feature space (red markers in far right panel; 50 draws were made from distributions for *A* and *B* and were compared to pseudo-randomly drawn inputs, black markers). When alternating the two learned or familiar patterns *A* and *B*, activations in the top layer of the networks experienced greater changes than when two randomly selected patterns were alternated (compare distances traversed in high-level feature space by red lines versus black lines, far right panel). Because of this large change in high-level activations, transient errors were generated back through the network for learned patterns. Strong oscillations could be observed in the error signals between stage 1 and stage 2 of the model for alternated familiar inputs *A* and *B* (120 ms period) (red curve, middle right panel). In contrast, alternating novel inputs with similar amplitude but with patterns not matching the learned weights led to small amplitude oscillations in upstream error signals (black curve). Note that the average response strength of state signals to novel and familiar inputs was matched at all network levels by construction (mean-matched inputs). The differing response dynamics of error signals under familiar versus novel patterns are qualitatively consistent with the IT findings for novel versus familiar images (left, reproduced with permission from *Figs. 1 & 2b* of Meyer et al., 2014). **(C)** The average response to the whole face for pIT sites recorded 0 to 1 mm below the pial surface (left panel; superficial recordings, red curve) and for sites 1 to 2.5 mm beneath the pial surface (black curve). (right panel) Average response of state units (red) and error units (black) in stage 1 of the model. Note the lagged response in state units which is similar to the lagged response of units in superficial recordings (red curve, left panel).

#### Evolution of neural signals across time

Neural responses to familiar images are known to rapidly attenuate in IT when compared to responses to novel images (Freedman, Riesenhuber, Poggio, & Miller, 2006)(Woloszyn & Sheinberg, 2012)(Meyer & Olson, 2011). This observation seems to contradict what would be predicted by simple Hebbian potentiation for the more exposed stimuli. Furthermore, familiar image responses show much sharper temporal dynamics than responses to novel images when presented repeatedly (Meyer et al., 2014). These qualitatively different dynamics for familiar versus novel images are surprising given that stimuli are drawn from the same distribution of natural images and are thus matched in their average image-level statistical properties (color, spatial frequency, contrast). To test whether our network displayed these different dynamical behaviors, we simulated familiar inputs as those that match the learned weight pattern of a high-level detector and novel inputs as those with the same overall input level but with weak correlation to the learned network weights (here, we have extended the network to include two units in the output stage corresponding to storage of the two familiarized input patterns to be alternated; conceptually, we consider these familiar pattern detectors as existing downstream of IT in a region such as perirhinal cortex which has been shown to code familiarized image statistics and memory-based object signals (Murray, Bussey, & Saksida, 2007)). We repeatedly alternated two familiar inputs or two novel inputs and found that error coding model responses in the hidden processing stage were temporally sharper for familiar inputs that matched the network’s feedforward weight patterns compared to novel patterns of input (pseudo-randomly drawn; see **Methods**), consistent with the previously observed phenomenon (**Figure 8B**; data reproduced with permission from Meyer et al., 2014). Model responses reproduced additional details of the neural dynamics including a large initial peak followed by smaller peaks for responses to novel inputs and a phase delay in the oscillations of responses to novel inputs compared to familiar inputs. Intuitively, these dynamics are composed of two phases after the initial response transient to the onset of the image sequence. In the first phase, familiar patterns lead to lower errors and hence lower neural responses than random patterns (**Figure 8B**, red curve drops below the black curve after the onset response), similar to the observed weaker response to more familiar face-like images present in our data (**Figure 2**, red curves drop below black curves in pIT example sites). When the familiar pattern *A* is switched to another familiar pattern *B*, this induces a short-term error in adjusting to the new pattern (**Figure 8B**, red curve briefly goes above the black curve during pattern switch and then decreases). In contrast, two unfamiliar patterns are closer together in the high-level encoding space than two learned patterns (**Figure 8B**, inset at right), and the switch between two unlearned patterns introduces relatively less shift in top-down signals and hence a smaller dynamical change in error signals. This result demonstrates that our model, derived from fitting only the first 70 ms (60-130 ms post image onset) of IT responses to face images, can extend to much longer timescales and may generalize to studies of images besides face images.

### Dynamical properties of neurons across cortical lamina

In the large family of state-error coding hierarchical networks, a number of different cortical circuits are possible. A key distinction of two such circuit mapping hypotheses (predictive coding versus error backpropagation) is the expected laminar location of state coding neurons transmitting information about features in the image. In typical neural network implementations, the feedforward projecting neurons in superficial lamina are presumed to encode estimates about states of the visual world (e.g. presence of a face). In contrast, predictive coding posits that superficial layers contain error units and that errors are projected forward to the next cortical level (Rao & Ballard, 1999)(Friston & Kiebel, 2009)(Hyvärinen, Hurri, & Hoyer, 2009). State and error signals can be distinguished by their dynamical signatures in our leading model, which was fit on error signals but produces predictions of the corresponding state signals underlying the generation of errors. Since state units are integrators (see **Methods**), they have slower dynamics than error units leading to longer response latencies and a milder decay in responses (**Figure 8C**, red curve in right panel). To test this prediction, we localized our recordings relative to the cortical mantle by co-registering the x-ray determined locations of our electrode (~400 micron *in vivo* accuracy; (Issa et al., 2010)) to structural MRI data (see **Methods**). When we separated units into those at superficial depths closer to the pial surface (1/3 of our sites; corresponds to approximately 0 to 1 mm in depth) versus those in the deeper layers (remaining 2/3 of sites, ~1 to 2.5 mm in depth), we found a longer latency and less response decay in superficial units consistent with the expected profile of state units (**Figure 8C**, left panel). Thus, the trend toward state-like signals in superficial layers is more consistent with typical error backpropagation models (states fed forward, errors fed backward) than with predictive coding proposals. In fact, the latency difference between cortical lamina in pIT (deep vs superficial: 66.0 ± 1.7 vs 76.0 ± 1.8 ms, p = 0.002) was greater than the conduction delay from pIT to cIT (i.e. from superficial layers of pIT to the deeper layers of cIT) (superficial pIT vs deep cIT: 76.0 ± 1.8 vs 75.5 ± 2.0 ms, p = 0.15) even though laminar distances within pIT are smaller than the distance traveled between cortical stages pIT and cIT. Thus, instead of a simple conduction delay accounting for latency differences across lamina, our model suggests that temporal integration of inputs in superficial lamina, more consistent with the behavior of state units as opposed to error units, may drive the lagged dynamical properties of neurons in superficial lamina.

### DISCUSSION

We have measured neural responses during a difficult frontal face detection task across the IT hierarchy and demonstrated that the population preference for faces in the intermediate (a.k.a hidden) processing stages decreases over time – that is population responses at lower levels of the hierarchy (pIT and cIT) rapidly evolved toward preferring unnatural, non-face part arrangements whereas the top level (aIT) rapidly developed and then maintained a preference for natural, frontal face part arrangements. The relative speed of selectivity changes in pIT (~30 ms) makes high-level explanations based on fixational eye movements or shifts in attention (e.g. from behavioral surprise to unnatural arrangements of face parts) unlikely as saccades and attention shifts occur on slower timescales (hundreds of milliseconds) (Egeth & Yantis, 1997). The presence of stronger responses to face images than to non-face images in aIT further argues against general arousal effects as these would have been expected to cause stronger responses to challenging non-face images in aIT. Rather, the rapid propagation of neural signals over tens of milliseconds suggested intracortical processing within the ventral visual stream in a manner that was not entirely consistent with a pure feedforward model, even when we included strong nonlinearities in these models such as normalization and even when we stacked these operations to form more complex three stage models. However, augmenting the feedforward model so that it represented the errors generated during hierarchical processing produced the observed neural dynamics and hierarchical signal propagation (**Figures 6-7**). This view argues that many IT neurons code error signals. Using this new modeling perspective, we went on to generate predictions of previously observed IT neural phenomena (**Figure 8**).

### Comparison to previous neurophysiology studies in IT

Our suggestion that many IT neurons code errors is consistent with the observation of strong responses to extremes in face space (Leopold, Bondar, & Giese, 2006) providing an alternative interpretation to the prior suggestion that cIT neurons are not tuned for typical faces but are instead tuned for atypical face features (i.e. extreme feature tuning) (Freiwald et al., 2009). In that prior work, the response preference of each neuron was determined by averaging over a long time window (~200 ms). By looking more closely at the fine time scale dynamics of the IT response, we suggest that this same extreme coding phenomenon can instead be interpreted as a natural consequence of networks that have an actual tuning preference for typical faces (as evidenced by an initial response preference for typical frontal faces in pIT, cIT, and aIT; **Figure 3B**) but that also compute error signals with respect to that preference. Under the present hypothesis, some IT neurons are preferentially tuned to typical spatial arrangements of face features, and other IT neurons are involved in coding errors with respect to those typical arrangements. We speculate that these intermixed state estimating and error coding neuron populations are both sampled in standard neural recordings of IT, even though only state estimating neurons are truly reflective of the tuning preferences of that IT processing stage. This intermixed view of IT neural signaling is further supported by recent studies demonstrating correlates of temporal prediction errors to image sequences in IT (Meyer & Olson, 2011) including the face-selective subregion of cIT (Schwiedrzik & Freiwald, 2017).

The precise fractional contribution of errors to total neural activity is difficult to estimate from our data. Under the primary image condition tested, not all sites significantly decreased their selectivity (~60% did not change their selectivity). We currently interpret these sites as coding state (feature) estimates (**Figure 3A**, light and dark gray lines in top and bottom rows, respectively), and we did observe evidence of emergence of state-like signals in our superficial neural recordings (**Figure 8C**). Alternatively, at least some of the non-reversing sites might be found to code errors under other image conditions than the one that we tested. Furthermore, while in our primary image condition selectivity decreases only accounted for ~15% of the overall spiking modulation (**Figure 6A**, data panel), larger modulations in late phase neural firing (50-100%) are possible under other image conditions (**Figure 8A,B**). At a computational level, the absolute contribution of error signals to spiking may not be the critical factor as even a small relative contribution may have important consequences in the network.

### Comparison across dynamical models of neural processing

Our goal was to test a range of existing recurrent models by recording neural dynamics across multiple cortical stages which provided stronger constraints on computational models than fitting neural responses from only one area as in prior work (Carandini et al., 1997)(Rao & Ballard, 1999). Crucially, we found that the multi-stage neural dynamics observed in our data could not be adequately fit by only using recurrences within a stage such as adaptation, lateral inhibition, and standard forms of normalization (**Figure 6**). These results did not change when we made our simple networks more complex by adding more stages (compare **Figure 6** versus **Figure 7**) or by using more realistic model units with monotonic nonlinearities similar to a spiking nonlinearity (data not shown). Indeed, we specifically chose our stimuli to evoke similar levels of within stage neural activity to limit the effects of known mechanisms that depend on activity levels within an area (e.g. adaptation, normalization), and we fully expect that these activity dependent mechanisms would operate in parallel to top-down, recurrent processes during general visual processing. We emphasize that we only tested the standard form of normalization as originally proposed, using within stage pooling and divisive mechanisms (Carandini et al., 1997). Since that original mechanistic formulation, normalization has evolved to become a term that broadly encapsulates many forms of suppression phenomena and can include both lateral interactions within an area and feedback interactions from other areas (Nassi, Gómez-Laberge, Kreiman, & Born, 2014)(Coen-Cagli, Kohn, & Schwartz, 2015). Thus, while our results do not follow from the original mechanistic form of normalization, they may yet fall under normalization more broadly construed as a term for suppression phenomena (error coding would require a similar suppressive component). Here, we have provided a normative model for how top-down suppression would follow from the well-defined computational goals of many hierarchical neural network models.

### Error signals generated across different hierarchical inference and learning models

The notion of error is inherent to many existing models in the literature that go beyond the basic feedforward, feature estimation class. Example models use errors for guiding top-down inference by computing errors implicitly (hierarchical Bayesian inference (Seung, 1997)(Lee & Mumford, 2003)) or by representing errors explicitly (predictive coding (Rao & Ballard, 1999)). In addition, errors can be used specifically for driving unsupervised learning (autoencoder (Rifai et al., 2011)) or supervised learning (classic error backpropagation (Williams & Hinton, 1986)). Recently, models have incorporated aspects of both inference and learning (Salakhutdinov & Hinton, 2012)(Patel, Nguyen, & Baraniuk, 2015). A key, unifying feature across inference and learning models is the need to compute an error signal between processing stages. This error signal can be in the form of a generative, reconstruction cost (stage *n* predicting stage *n-1*) or a discriminative, construction cost (stage *n-1* predicting stage *n*). Across-stage “performance” error terms are used in all model cost functions, are typically the only term combining signals from different model stages, and are distinct from within-stage “regularization” terms (i.e. sparseness or weight decay) in driving network behavior (Marblestone, Wayne, & Kording, 2016). The present study provides evidence that such errors are not only computed, but that they are explicitly encoded in spiking rates. We emphasize that this result at the level of population dynamics was robust across choices of cost function such as those used in the literature; we tested models with different unsupervised and supervised performance errors (reconstruction, nonlinear reconstruction, and discriminative) and found similar population level error signals across these networks in the basic two-layer implementation (**Figure 7C**). Thus, errors as generally instantiated in the state-error coding hierarchical model family provide a good approximation to IT population neural dynamics.

### Computational utility of coding errors in addition to states

In error-computing networks, errors provide control signals for guiding learning giving these networks additional adaptive power over basic feature estimation networks. This property helps augment the classical, feature coding view of neurons which, with only feature activations and Hebbian operations, does not lead to efficient learning in the manner produced by gradient descent using error backpropagation (Williams & Hinton, 1986). Observation of error signals may provide insight into how more intelligent unsupervised and supervised learning algorithms such as backpropagation could be plausibly implemented in the brain. A potentially important contribution of this work is the suggestion that gradient descent algorithms are facilitated by using an error code so that efficient learning is reduced to a simple Hebbian operation at synapses and efficient inference is simply integration of inputs at the cell body (see *eqn 10* and text in **Methods**). This representational choice, to code the computational primitives of gradient descent in spiking activity, would simply leverage the existing biophysical machinery of neurons for inference and learning.

## MATERIALS & METHODS

### Animals and surgery

All surgery, behavioral training, imaging, and neurophysiological techniques are identical to those described in detail in previous work (Issa & DiCarlo, 2012). Two rhesus macaque monkeys (*Macaca mulatta*) weighing 6 kg (Monkey 1, female) and 7 kg (Monkey 2, male) were used. A surgery using sterile technique was performed to implant a plastic fMRI compatible headpost prior to behavioral training and scanning. Following scanning, a second surgery was performed to implant a plastic chamber positioned to allow targeting of physiological recordings to posterior, middle, and anterior face patches in both animals. All procedures were performed in compliance with National Institutes of Health guidelines and the standards of the MIT Committee on Animal Care and the American Physiological Society.

### Behavioral training and image presentation

Subjects were trained to fixate a central white fixation dot during serial visual presentation of images at a natural saccade-driven rate (one image every 200 ms). Although a 4° fixation window was enforced, subjects generally fixated a much smaller region of the image (<1°) (Issa & DiCarlo, 2012). Images were presented at a size of 6° except for control tests at 3° and 12° sizes (**Figure 4C**), and all images were presented for 100 ms duration with 100 ms gap (background gray screen) between each image. Up to 15 images were presented during a single fixation trial, and the first image presentation in each trial was discarded from later analyses. Five repetitions of each image in the general screen set were presented, and ten repetitions of each image were collected for all other image sets. The screen set consisted of a total of 40 images drawn from four categories (faces, bodies, objects, and places; 10 exemplars each) and was used to derive a measure of face versus non-face object selectivity. Following the screen set testing, some sites were tested using an image set containing images of face parts presented in different combinations and positions (**Figure 1B**, left panel). We first segmented the face parts (eye, nose, mouth) from a monkey face image. These parts were then blended using a Gaussian window, and the face outline was filled with pink noise to create a continuous background texture. A face part could appear on the outline at any one of nine positions on an evenly spaced 3×3 grid. Although the number of possible images is large (4^9^ = 262,144 images), we chose a subset of these images for testing neural sites (n = 82 images). Specifically, we tested the following images: the original whole face image, the noise-filled outline, the whole face reconstructed by blending the four face parts with the outline, all possible single part images where the eye, nose, or mouth could be at one of nine positions on the outline (n = 3×9 = 27 images), all two part images containing a nose, mouth, left eye, or right eye at the correct outline-centered position and an eye tested at all remaining positions (n = 4*8-1 = 31 images), all two part images containing a correctly positioned contralateral eye while placing the nose or mouth at all other positions (n = 2*8-2 = 14 images), and all correctly configured faces but with one or two parts missing besides those already counted above (n = 4+3 = 7 images). The particular two-part combinations tested were motivated by prior work demonstrating the importance of the eye in early face processing (Issa & DiCarlo, 2012), and we sought to determine how the position of the eye relative to the outline and other face parts was encoded in neural responses. The three and four part combinations were designed to manipulate the presence or absence of a face part for testing the integration of face parts, and in these images, we did not vary the positions of the parts from those in a naturally occurring face. In a follow-up test on a subset of sites, we permuted the position of the four face parts under the constraint that they still formed the configuration of a naturally occurring face (i.e. preserve the ‘T’ configuration, n = 10 images; **Figure 4B**). We tested single part images at 3° and 12° sizes in a subset of sites (n = 27 images at each size; **Figure 4C**). Finally, we measured the responses to the individual face parts in the absence of the outline (n = 4 images; **Figure 8A**).

### MR Imaging and neurophysiological recordings

Both structural and functional MRI scans were collected in each monkey. Putative face patches were identified in fMRI maps of face versus non-face object selectivity in each subject. A stereo microfocal x-ray system (Cox et al., 2008) was used to guide electrode penetrations in and around the fMRI defined face-selective subregions of IT. X-ray based electrode localization was critical for making laminar assignments since electrode penetrations are often not perpendicular to the cortical lamina when taking a dorsal-ventral approach to IT face patches. Laminar assignments of recordings were made by co-registering x-ray determined electrode coordinates to MRI where the pial-to-gray matter border and the gray-to-white matter border were defined. Based on our prior work estimating sources of error (e.g. error from electrode tip localization and brain movement), registration of electrode tip locations to MRI brain volumes has a total of <400 micron error which is sufficient to distinguish deep from superficial layers (Issa et al., 2013). Multi-unit activity (MUA) was systematically recorded at 300 micron intervals starting from penetration of the superior temporal sulcus such that all sites at these regular intervals were tested with a screen set containing both faces and non-face objects, and a subset of sites that were visually driven were further tested with our main image set manipulating the position of face parts. Although we did not record single-unit activity, our previous work showed similar responses between single-units and multi-units on images of the type presented here (Issa & DiCarlo, 2012), and our results are consistent with observations in previous single-unit work in IT (Freiwald et al., 2009). Recordings were made from PL, ML, and AM in the left hemisphere of monkeys 1 and 2 and additionally from AL in monkey 2. AM and AL are pooled together in our analyses forming the aIT sample while PL and ML correspond to the pIT and cIT samples, respectively.

### Neural data analysis

The face patches were physiologically defined in the same manner as in our previous study (Issa & DiCarlo, 2012). Briefly, we fit a graded 3D sphere model (linear profile of selectivity that rises from a baseline value toward the maximum at the center of the sphere) to the spatial profile of face versus non-face object selectivity across our sites. We tested spherical regions with radii from 1.5 to 10 mm and center positions within a 5 mm radius of the fMRI-based centers of the face patches. The resulting physiologically defined regions were 1.5 to 3 mm in diameter. Sites which passed a visual response screen (mean response in a 60-160 ms window >2*SEM above baseline for at least one of the four categories in the screen set) were included in further analysis. All firing rates were baseline subtracted using the activity in a 25-50 ms window following image onset averaged across all repetitions of an image. Finally, given that the visual response latencies in monkey 2 were on average 13 ms slower than those in monkey 1 for corresponding face-selective regions, we applied a single latency correction (13 ms shift to align monkey 1 and monkey 2’s data) prior to averaging across monkeys. This was done so as not to wash out any fine timescale dynamics by averaging. Similar results were obtained without using this latency correction as dynamics occurred at longer timescales (~30 ms). This single absolute adjustment was more straightforward than the site-by-site adjustment used in our previous work (Issa & DiCarlo, 2012) (though similar results were obtained using this alternative latency correction); even when each monkey was analyzed separately, we still observed pIT selectivity dynamics (**Figure 4A**). Images that produced an average population response ≥ 0.9 of the initial response (60-100 ms) to a face image with all face parts arranged in their typical positions in a frontal face were analyzed further (**Figures 2 and 3**). Stimulus selection was intended to limit potentially confounding differences in visual drive between image classes. In a control test, we also repeated our analysis by selecting images on a site-by-site basis where images with frontal face and non-face arrangements of parts were chosen to be within 0.75x to 1.25x of the initial response to the complete face image (minimum of five face and five non-face images in this response range for inclusion of site in analysis). In follow-up analyses of population responses, we specifically limited comparison to images with the same number of parts (**Figure 4B,C**). For example, for single part images, we used the image with the eye in the upper, contralateral region of the outline as a reference requiring a response ≥ 0.9 of the initial population response to this reference for inclusion of the images in this analysis. We found that four other images of the 27 single-part images elicited a response at least as large as 90% of the response to this standard image. For images containing all four face parts, we used the complete, frontal face as the standard and found non-face arrangements of the four face parts that drove at least 90% of the early response to the whole face (2 images out of 10 tested). To compute individual site d’ for each of these stimulus partitions (e.g. typical versus atypical arrangements of 1 face part), we combined all presentations of images with frontal face arrangements and compared these responses to responses from all presentations of images with non-face arrangements using *d’ =* (*u_1_- u_2_*)/((*var*_*1*_+*var*_*2*_)/2)^1/2^ where variance was computed across all trials for that image class (e.g. all presentations of all typical face images); this was identical to the d’ measure used in previous work for computing selectivity for faces versus non-face objects (Aparicio, Issa, & DiCarlo, 2016; Ohayon, Freiwald, & Tsao, 2012). For example, for the main image set (**Figure 2A**), we compared all presentations of frontal face arrangements (8 images x 10 presentations/image = 80 total presentations) to all presentations of non-face arrangements (13 images x 10 presentations/image = 130 total presentations) to compute the d’ values for each site in two time windows (60-90 ms and 100-130 ms) as shown in **Figure 3A**. A positive d’ implies a stronger response to more naturally occurring frontal face arrangements of face parts while a negative d’ indicates a preference for unnatural non-face arrangements of the face parts.

## Dynamical models

### Modeling framework and equations

To model the dynamics of neural response rates in a hierarchy, we start with the simplest possible model that might capture those dynamics: a model architecture consisting of a hidden stage of processing containing two units that linearly converge onto a single output unit. We use this two-stage cascade for illustration of the basic concepts which can be easily extended to longer cascades with additional stages, and we ultimately used a three-stage version of the model to fit our neural data collected from three cortical stages (**Figures 5B & 6**).

An external input is applied separately to each hidden stage unit, which can be viewed as representing different features for downstream integration. We vary the connections between the two hidden units within the hidden processing stage (lateral connections) or between hidden and output stage units (feedforward and feedback connections) to instantiate different model families. The details of the different architectures specified by each model class can be visualized by their equivalent neural network diagrams (**Figure 5**). Here, we provide a basic description for each model tested. All two-stage models utilized a 2×2 feedforward identity matrix *A* that simply transfers inputs ***u*** (2×1) to hidden layer units ***x*** (2×1) and a 1×2 feedforward vector *B* that integrates hidden layer activations ***x*** into a single output unit *y*.

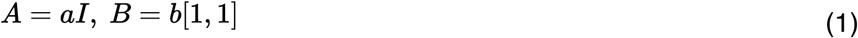

By simply substituting in the appropriate unit vector and weight matrix transforming inputs from one layer to the next for the desired network architecture, this simple two-stage architecture can be extended to larger networks (e.g. see three-stage network diagrams in **Figure 5B**). To generate dynamics in the simple networks below, we assumed that neurons act as leaky integrators of their total synaptic input, a standard rate-based model of a neuron used in previous work (Seung, 1997)’(Rao & Ballard, 1999).

### Pure feedforward

In the purely feedforward family, connections are exclusively from hidden to output stages through feedforward matrices *A* and *B*.

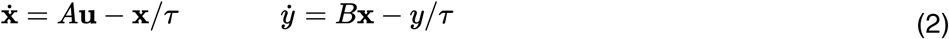

where *τ* is the time constant of the leak current which can be seen as reflecting the biophysical limitations of neurons (a perfect integrator with large *τ* would have almost no leak and hence infinite memory).

### Lateral inhibition

Lateral connections (matrix with off-diagonal terms) are included and are inhibitory. The scalar *k*_*l*_ sets the relative strength of lateral inhibition versus bottom-up input.

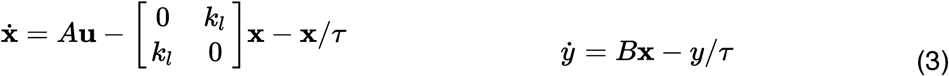

*Normalization*. An inhibitory term that scales with the summed activity of units within a stage is included. The scalar *k*_*s*_ sets the relative strength of normalization versus bottom-up input.

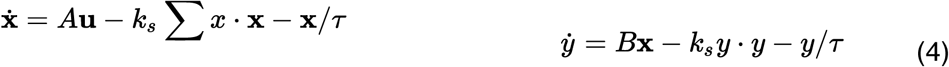

*Normalization (nonlinear)* (Carandini et al., 1997). The summed activity of units within a stage is used to nonlinearly scale shunting inhibition.

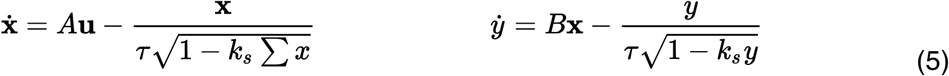

Note that this is technically a nonlinear dynamical system, and since the normalization term in equation (5) is not continuously differentiable, we used the fourth-order Taylor approximation around zero in the simulations of equation (5).

### Feedback (linear reconstruction)

The feedback-based model is derived using a normative framework that performs optimal inference in the linear case (Seung, 1997) (unlike the networks in equations (2)-(5) which are motivated from a mechanistic perspective but do not directly optimize a squared error performance loss). The feedback network minimizes the cost *C* of reconstructing the inputs of each stage (i.e. mean squared error of layer *n* predicting layer *n-1*).

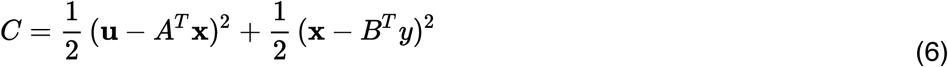

Differentiating this coding cost with respect to the encoding variables in each layer ***x***, *y* yields:

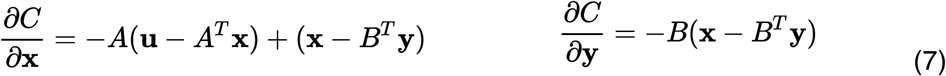

The cost function *C* can be minimized by descending these gradients over time to optimize the values of ***x*** and *y*:

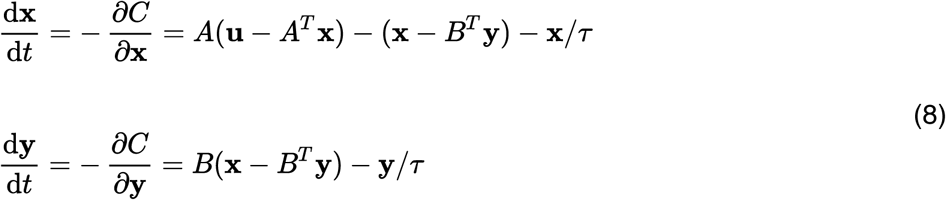

The above dynamical equations are equivalent to a linear network with a connection matrix containing symmetric feedforward (*B*) and feedback (*B*^*T*^) weights between stages ***x*** and *y* as well as within-stage pooling followed by recurrent inhibition (-*AA^T^**x*** and *-BB^T^y*) that resembles normalization. The property that symmetric connections minimize the cost function *C* generalizes to a feedforward network of any size or number of hidden processing stages (i.e. holds for arbitrary lower triangular network connection matrix). The final activation states (***x***,*y*) of the hierarchical generative network are optimal in the sense that the bottom-up activations (implemented through feedforward connections) are balanced by the top-down expectations (implemented by feedback connections) which is equivalent to a Bayesian network combining bottom-up likelihoods with top-down priors to compute the maximum *a posteriori* (MAP) estimate. Here, the priors are embedded in the weight structure of the network. In simulations, we include an additional scalar *k*_*td*_ that sets the relative weighting of bottom-up versus top-down signals.

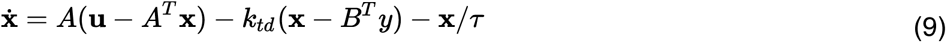

### Error signals computed in the feedback model

In equation (9), inference can be thought of as proceeding through integration of inputs on the dendrites of neuron population ***x***. In this scenario, all computations are implicit in dendritic integration. Alternatively, the computations in equation (9) can be done in two steps where, in the first step, reconstruction errors are computed (i.e. ***e_0_** = **u**-A^T^**x**, **e_1_** = **x**-B^T^y*) and explicitly represented in a separate error coding population. These error signals can then be integrated by their downstream target population to generate the requisite update to the state signal of neuron population ***x***.

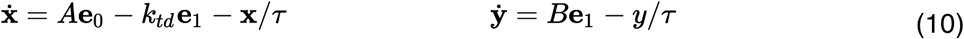

An advantage of this strategy is that the a state unit now directly receives errors as inputs, and those inputs allow implementation of an efficient Hebbian rule for learning weight matrices (Rao & Ballard, 1999) -- the gradient rule for learning is simply a product of the state activation and the input error activation (weight updates obtained by differentiating equation (6) with respect to weight matrices *A* and *B*: 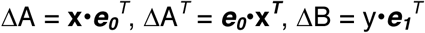, and 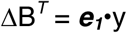. Thus, the reconstruction errors serve as computational intermediates for both the gradients of online inference mediated by dendritic integration (dynamics in state space, equation (10)) and gradients for offline learning mediated by Hebbian plasticity (dynamics in weight space).

In order for the reconstruction errors at each layer to be scaled appropriately in the feedback model, we invoke an additional downstream variable *z* to predict activity at the top stage such that, instead of ***e_2_** = y* which scales as a state variable, we have ***e_2_** = y-C^T^z* (**Figure 5A**). This overall model reflects a state and error coding model as opposed to a state only model.

### Feedback (three-stage)

For the simulations in **Figures 6 and 8**, three-stage versions of the above equations were used. These deeper networks were also wider such that they began with four input units (***u***) instead of only two inputs in the two-stage models. These inputs converged through successive processing stages (***w,x,y***) to one unit at the top node (***z***) (**Figure 5B**).

### Feedback (nonlinear reconstruction)

To test the generality of our findings beyond a linear reconstruction cost, we simulated feedback-based models which optimized different candidate cost functions proposed for the ventral stream (**Figure 7C**). In nonlinear hierarchical inference, reconstruction is performed using a monotonic nonlinearity with a threshold (*th*) and bias (*bi*):

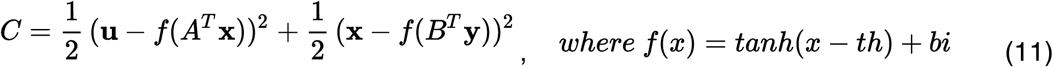

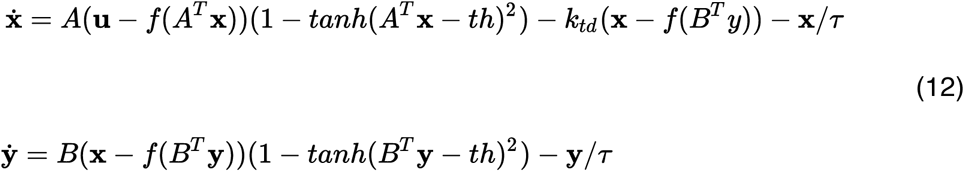

### Feedback (linear construction)

Instead of a reconstruction cost where responses match the input (i.e. generative model) as in unsupervised learning, we additionally simulated the states and errors in a feedback network minimizing a linear construction cost where the network is producing responses to match a given output (i.e. discriminative model) similar to supervised learning:

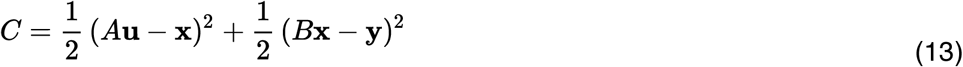

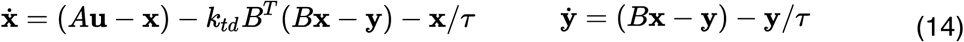

### Model simulation

To simulate the dynamical systems in equations (2)-(14), a step input ***u*** was applied. This input was smoothed using a Gaussian kernel to approximate the lowpass nature of signal propagation in the series of processing stages from the retina to pIT:

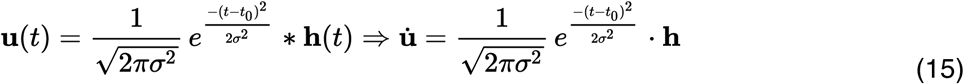

where the elements of ***h*** are scaled Heaviside step functions. The input is thus a sigmoidal ramp whose latency to half height is set by *t*_*0*_ and rise time is set by σ. For simulation of two-stage models, there were ten basic parameters: latency of the input *t*_*0*_, standard deviation of the Gaussian ramp *σ*, system time constant *τ*, input connection strength *a*, feedforward connection strength *b*, the four input values across two stimulus conditions (i.e. *h*_*11*_, *h*_*12*_, *h*_*21*_, *h*_*22*_), and a factor *sc* for scaling the final output to the neural activity. In the deeper three-stage network, there were a total of fifteen parameters which included an additional feedforward connection strength *c* and additional input values since the three-stage model had four inputs instead of two. The lateral inhibition model class required one additional parameter *k*_*l*_ as did the normalization model family *k*_*s*_, and for feedback model simulations, there was an additional feedback weight *k*_*td*_ to scale the relative contribution of the top-down errors in driving online inference. For the error coding variants of the feedback model, gain parameters *c* (two-stage) and *d* (three-stage) were included to scale the overall magnitude of the top level reconstruction error (also see **Figure 5** for locations of parameters in network diagrams).

### Model parameter fits to neural data

In fitting the models to the observed neural dynamics, we mapped the summed activity in the hidden stage (***w***) to population averaged activity in pIT, and we mapped the summed activity in the output stage (***y***) to population averaged signals measured in aIT. To simulate error coding, we mapped the reconstruction errors ***e_1_** = **w**-B^T^**x*** and ***e_3_** = y-C^T^z* to activity in pIT and aIT, respectively. We applied a squaring nonlinearity to the model outputs as an approximation to rectification since recorded extracellular firing rates are non-negative (and linear rectification is not continuously differentiable). Analytically solving this system of dynamical equations (2)-(14) for a step input is precluded because of the higher order interaction terms (the roots of the determinant and hence the eigenvalues/eigenvectors of a 3×3 or larger matrix are not analytically determined, except for the purely feedforward model which only has first-order interactions), and in the case of the normalization models, there is an additional nonlinear dependence on the shunt term. Thus, we relied on computational methods (constrained nonlinear optimization) to fit the parameters of the dynamical systems to the neural data with a quadratic (sum of squares) loss function.

Parameter values were fit in a two-step procedure. In the first step, we fit only the difference in response between image classes (differential mode which is the selectivity profile over time, see **Figure 6A**, right data panel), and in the second step, we refined fits to capture an equally weighted average of the differential mode and the common mode (the common mode is the average across images of the response time course of visual drive). This two-step procedure was used to ensure that each model had the best chance of fitting the dynamics of selectivity (differential mode) as these selectivity profiles were the main phenomena of interest but were smaller in size (20% of response) compared to overall visual drive. In each step, fits were done using a large-scale algorithm (interior-point) to optimize coarsely, and the resulting solution was used as the initial condition for a medium-scale algorithm (sequential quadratic programming) for additional refinement. The lower and upper parameter bounds tested were: *t*_*0*_=[50 70], σ=[0.5 25], τ=[0.5 1000], *k*_*l*_,*k*_*s*_,*k*_*td*_=[0 1], *a,b*,*c,d*=[0 2], *h*=[0 20], *sc*=[0 100], *th*=[- 20 20], and *bi*=[-1 1] which proved to be adequately liberal as parameter values converged to values that did not generally approach these boundaries. To avoid local minima, the algorithm was initialized to a number of randomly selected points (n = 50), and after fitting the differential mode, we took the top fits (n = 25) for each model class and used these as initializations in subsequent steps. The single best fitting instance of each model class is shown in the main figures.

### Model predictions

For the predictions in **Figure 8**, all architectural parameters obtained by the fitting procedure above were held fixed; only the pattern of inputs to the network was varied. For **Figure 8A**, to test the input integration properties of a model, we used the top-performing model and compared the response to all inputs presented simultaneously with the sum of the responses to each input alone.

For **Figure 8B**, we approximated novel versus familiar images by parametrically varying the degree to which input patterns were random (inputs drawn from i.i.d. uniform distributions on the interval [0,1]) versus structured in a way that matched the weight pattern of the network. Here, we used a version of the model with two independent outputs reflecting detectors for two familiarized input patterns (output 1 tuned to pattern *A*: *u*_*1*_, *u*_*2*_, *u*_*3*_, *u*_*4*_ active and output 2 tuned to pattern 2: *u*_*5*_, *u*_*6*_, *u*_*7*_, *u*_*8*_ active) (**Figure 8B**). Alternating between these two input patterns simulates alternation of two familiarized (learned) images as compared to purely random patterns (*u*_*1-8*_ independent and identically distributed). The first-to-second layer weights were [1,1,1,1,0,0,0,0] for pattern A and [0,0,0,0,1,1,1,1] for pattern B, so to parametrically vary the degree of correlation of inputs to this weight pattern from random (correlation = 0) to deterministic (correlation = 1), we drew input values from a joint distribution *P(u_1_*,*u*_*2*_,*u*_*3*_,*u*_*4*_,*u*_*5*_,*u*_*6*_,*u*_*7*_,*u*_*8*_) where *u*_*1-4*_ were drawn from a high-valued uniform distribution on the interval [1-ɛ,1] and *u*_*5-8*_ were drawn from a low-valued uniform distribution [0, ε] for stimulus pattern *A* and the opposite for pattern *B* (*u*_*5-8*_ high-valued and *u*_*1-4*_ low-valued). The parameter ε determines the range of values that could be drawn from purely deterministic (0 or 1, ɛ = 0) to randomly uniformly distributed (from 0 to 1, ɛ = 1). Thus, the correlation of the inputs correspondingly varies according to ρ(*u*_*i*_,*u*_*j*_) = ρ(*u*_*k*_,*u*_*l*_) = (ɛ-1)^2^/((ɛ-1)^2^+ɛ^2^/3) where 1≤*i*,*j*≤4, *i*≠*j* and 5≤*k*,*l*≤8, *k*≠*l* approaching correlation equal to 0 for a purely, random pattern (ε= 1) that had a low probability of matching the learned patterns *A* and *B*.

### Code availability

All data analysis and computational modeling were done using custom scripts written in Matlab. All code is available upon request.

### Statistics

Error bars represent standard errors of the mean obtained by bootstrap resampling (n = 1000). All statistical comparisons including those of means or correlation values were obtained by bootstrap resampling (n = 1000) producing p-values at a resolution of 0.001 so that the lowest p-value that can be reported is p = 0.000 given the resolution of this statistical analysis. All statistical tests were two-sided unless otherwise specified. Spearman’s rank correlation coefficient was used.

## ACKNOWLEDGEMENTS

We thank J. Deutsch, K. Schmidt, and P. Aparicio for help with MRI and animal care and B. Andken and C. Stawarz for help with experiment software. We are grateful to T. Meyer and C. Olson for sharing figures from their published work.

